# Does higher sampling rate (Multiband + SENSE) benefit the detection of task correlated BOLD for cognitive neuroscience applications at 3T?

**DOI:** 10.1101/762831

**Authors:** Ritu Bhandari, Evgeniya Kirilina, Matthan Caan, Judith Suttrup, Teresa de Sanctis, Lorenzo De Angelis, Christian Keysers, Valeria Gazzola

**Author notes:** Equally contributing second authors (Evgeniya Kirilina, Matthan Caan). Equally contributing third authors (Judith Suttrup, Teresa de Sanctis, Lorenzo De Angelis). Equally contributing last authors (Christian Keysers, Valeria Gazzola).

## Abstract

Multiband (MB) or Simultaneous multi-slice (SMS) acquisition schemes allow the acquisition of MRI signals from more than one spatial coordinate at a time. Commercial availability has brought this technique within the reach of many neuroscientists and psychologists. Most early evaluation of the performance of MB acquisition employed resting state fMRI or the most basic tasks. In this study, we tested whether the advantages of using MB acquisition schemes generalize to group analyses using a cognitive task more representative of typical cognitive neuroscience applications. Twenty-three subjects were scanned on a Philips 3T scanner using five sequences up to eight-fold acceleration with MB-factors 1 to 4, SENSE factors up to 2 and corresponding TRs of 2.45s down to 0.63s, while they viewed (i) movies showing complex actions with hand object interactions and (ii) control movies without hand object interaction. Using random effects group-level, voxel-wise analysis we found that all sequences were able to detect the basic action observation network known to be recruited by our task. The highest t-values were found for sequences with MB4 acceleration. For the MB1 sequence, a 50% bigger voxel volume was needed to reach comparable t-statistics. The group-level t-values for resting state networks (RSNs) were also highest for MB4 sequences. Here the MB1 sequence with larger voxel size did not perform comparable to the MB4 sequence. Altogether, we can thus recommend the use of MB4 (and SENSE 1.5 or 2) on a Philips scanner when aiming to perform group-level analyses using cognitive block design fMRI tasks and voxel sizes in the range of cortical thickness (e.g. 2.7mm isotropic). While results will not be dramatically changed by the use of multiband, our results suggest that MB will bring a moderate but significant benefit.

## 1 Introduction

Multiband (MB) or Simultaneous multi-slice (SMS) acquisition schemes allow the acquisition of magnetic resonance imaging (MRI) signals from more than one spatial coordinate at a time. Apart from the obvious advantage in the reduction of per volume acquisition times (Moeller et al., 2010), these sequences have been postulated to have several advantages for functional MRI (fMRI), especially, higher signal to noise per time unit, higher sampling rate resulting in higher statistical power and a better estimation of physiological noise (Barth, Breuer, Koopmans, Norris, & Poser, 2016; Feinberg & Setsompop, 2013; Olafsson, Kundu, Wong, Bandettini, & Liu, 2015). Most of the early testing of performance of MB acquisition employed resting state fMRI and showed decreased total scan times (X.-H. Liao et al., 2013), better noise estimation and spectral and spatial de-aliasing (Griffanti et al., 2014; Kalcher et al., 2014; Tong & Frederick, 2014a, 2014b; Tong, Hocke, & Frederick, 2014) as well as better localization and estimation of functional networks (Koopmans, Boyacioğlu, Barth, & Norris, 2012). However, much less information is available about the benefits of MB acquisition for task based fMRI (Feinberg & Yacoub, 2012). Particularly, the utility of MB acquisition for group level statistics has not been evaluated yet. In this study we therefore performed comprehensive analyses to test the advantages of using MB acquisition schemes over single band EPI acquisition schemes in detecting task-based Blood Oxygenation Level Dependent (BOLD) responses.

The idea of acquiring data from different spatial locations simultaneously was first proposed in the 1980s (see Barth et al., 2016 for a historical review). However, the studies that brought widespread attention to the use of MB technique in fMRI were published in 2010 as part of the Human Connectome Project (Smith et al., 2013; Uğurbil et al., 2013). These two studies used multiplexed EPI combining simultaneous echo refocusing (SIR) and multiband radio frequency pulses and showed increased sensitivity to detect resting state networks (RSN) with 6 fold higher sampling rate at 3 Tesla (T) (Feinberg et al., 2010), and comparable activation between single band and MB acquisition using simple sensory tasks at 7T (Moeller et al., 2010). Since then, MB sequences were improved to include multiple modalities (Cohen, Nencka, Lebel, & Wang, 2017), to reduce radiofrequency power deposition (Auerbach, Xu, Yacoub, Moeller, & Uğurbil, 2013; Koopmans et al., 2012; Norris, Koopmans, Boyacioğlu, & Barth, 2011; Wu et al., 2013) and g-factor penalty due to the suboptimal reconstruction of the images (Setsompop et al., 2012), to ameliorate image reconstruction using controlled aliasing methods (Blaimer, Choli, Jakob, Griswold, & Breuer, 2013) such as radial CAIPIRINHA (Yutzy, Seiberlich, Duerk, & Griswold, 2011) and blipped CAIPI techniques (Setsompop et al., 2012), and by optimizing coils design (Poser et al., 2014). Together with these technical advancements in MR pulse sequence development and image quality improvements, the commercial availability of compatible hardware and tailored acquisition and reconstruction software has brought the technique within the reach of many neuroscientists and psychologists, paving the way to systematic testing, validation and application to basic and clinical research settings (Chekroud, Anand, Yong, Pike, & Bridge, 2017; Cohen-Gilbert et al., 2017; Gabrielsen et al., 2018; Harenski et al., 2018; Kim, Zhao, & Bae, 2016; King et al., 2018; Kyathanahally, Wang, Calhoun, & Deshpande, 2017; Provencher, Bizeau, Gilbert, Bérubé-Lauzière, & Whittingstall, 2018; Shah, Cramer, Ferguson, Birn, & Anderson, 2016; Suri et al., 2017).

Since the publication of the first fMRI studies using MB acquisition in 2010 (Feinberg et al., 2010; Moeller et al., 2010), several studies have tested its benefits on resting and task based functional data. Using the keywords ‘Multiband EPI’, ‘Multiband fMRI’, ‘Simultaneous Multislice EPI’ and ‘Simultaneous Multislice fMRI’ in Pubmed on 15 July 2019, we identified 40 publications (including the two 2010 publications) that assessed the advantages of multiband acquisition for studying resting state and/or task based fMRI (Table1). The data presented in these studies was collected on either 3T or 7T field strength scanners, generally using small to moderate sample sizes (median= 10; range = 3 to 476), and different voxel sizes (ranging between 1.5 mm isotropic and 3.75 mm isotropic). Of these 40 studies, 20 studies used resting state acquisition not only to assess the performance of low repetition times (TRs) (Feinberg et al., 2010; Koopmans et al., 2012; X.-H. Liao et al., 2013; Preibisch, Castrillón G., Bührer, & Riedl, 2015) but also to harness the advantages of the low TRs in spectral de-aliasing of high frequency bands to study the BOLD specific as well as noise specific components in different frequency bands, and for ICA based de-noising methods (Boubela et al., 2013; Gohel & Biswal, 2015; Griffanti et al., 2014; Kalcher et al., 2014; Tong & Frederick, 2014a).

**Table1.**
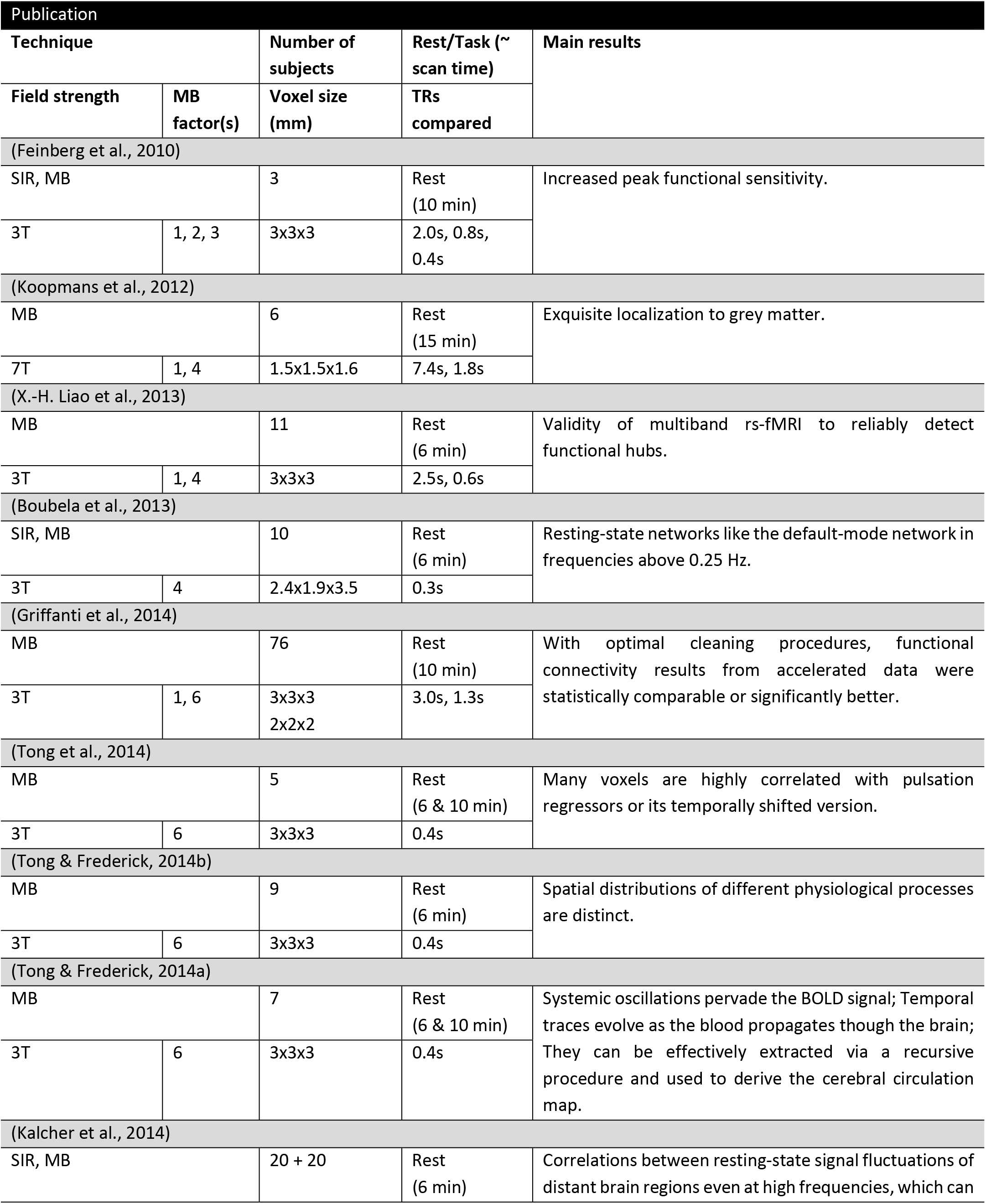
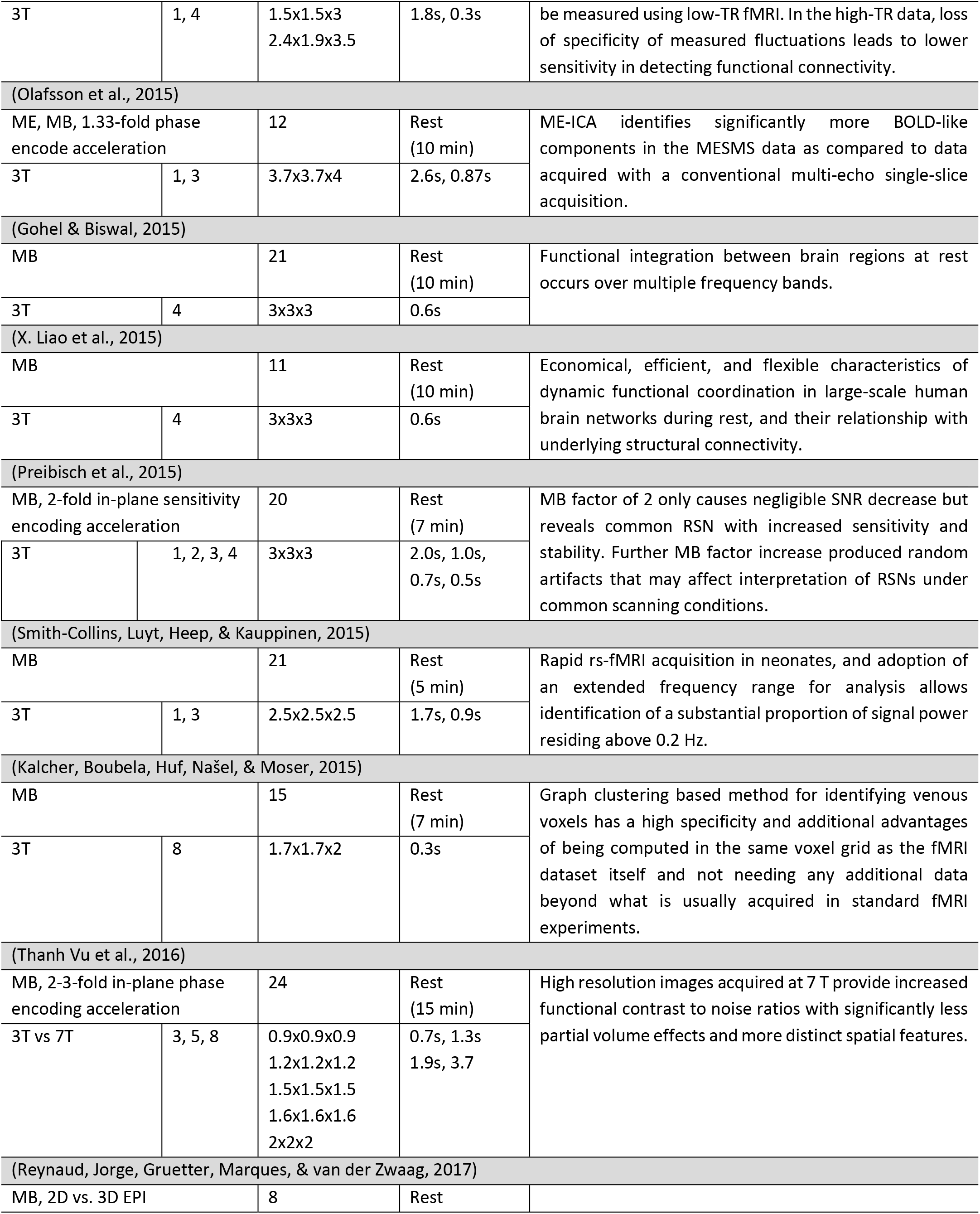
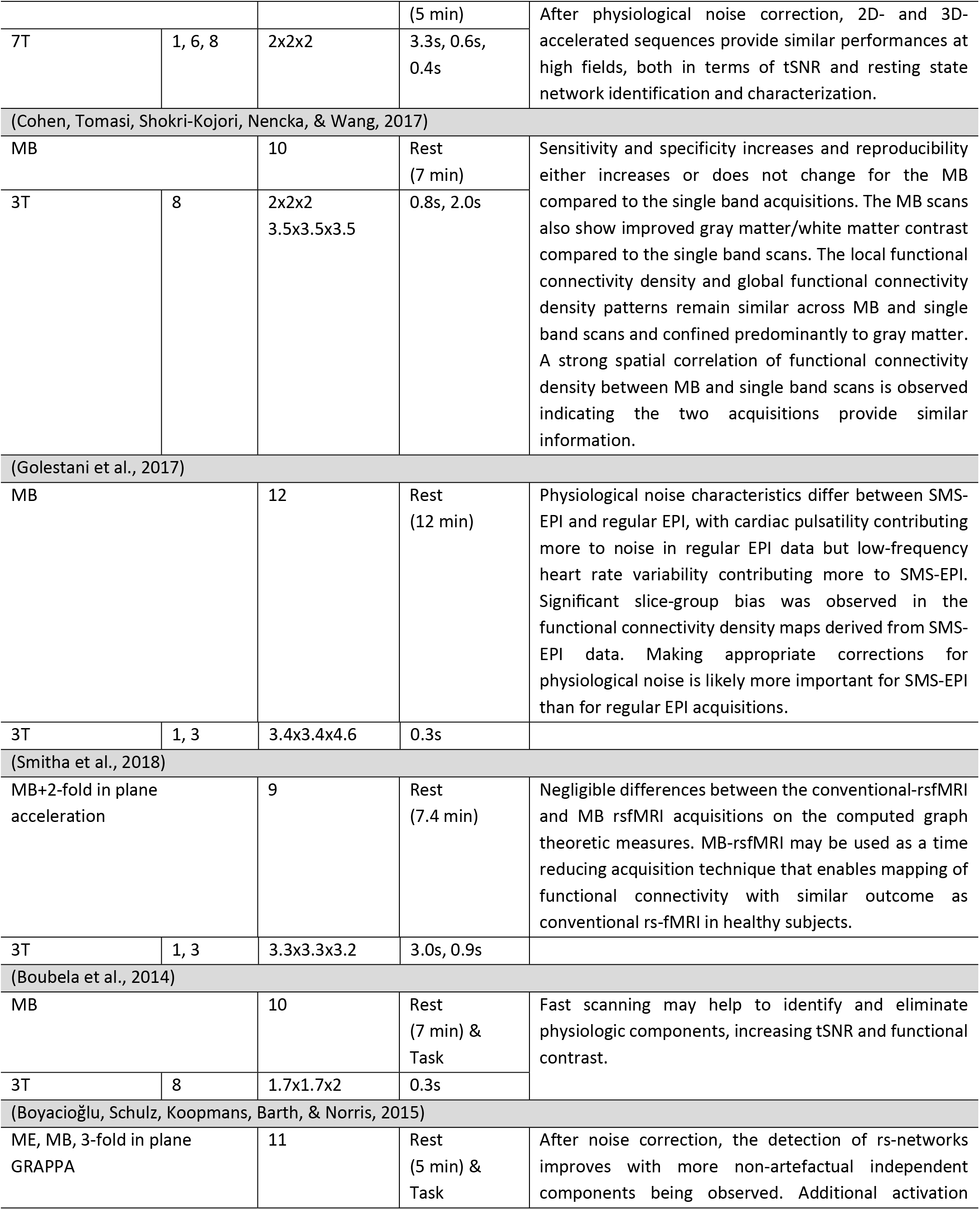
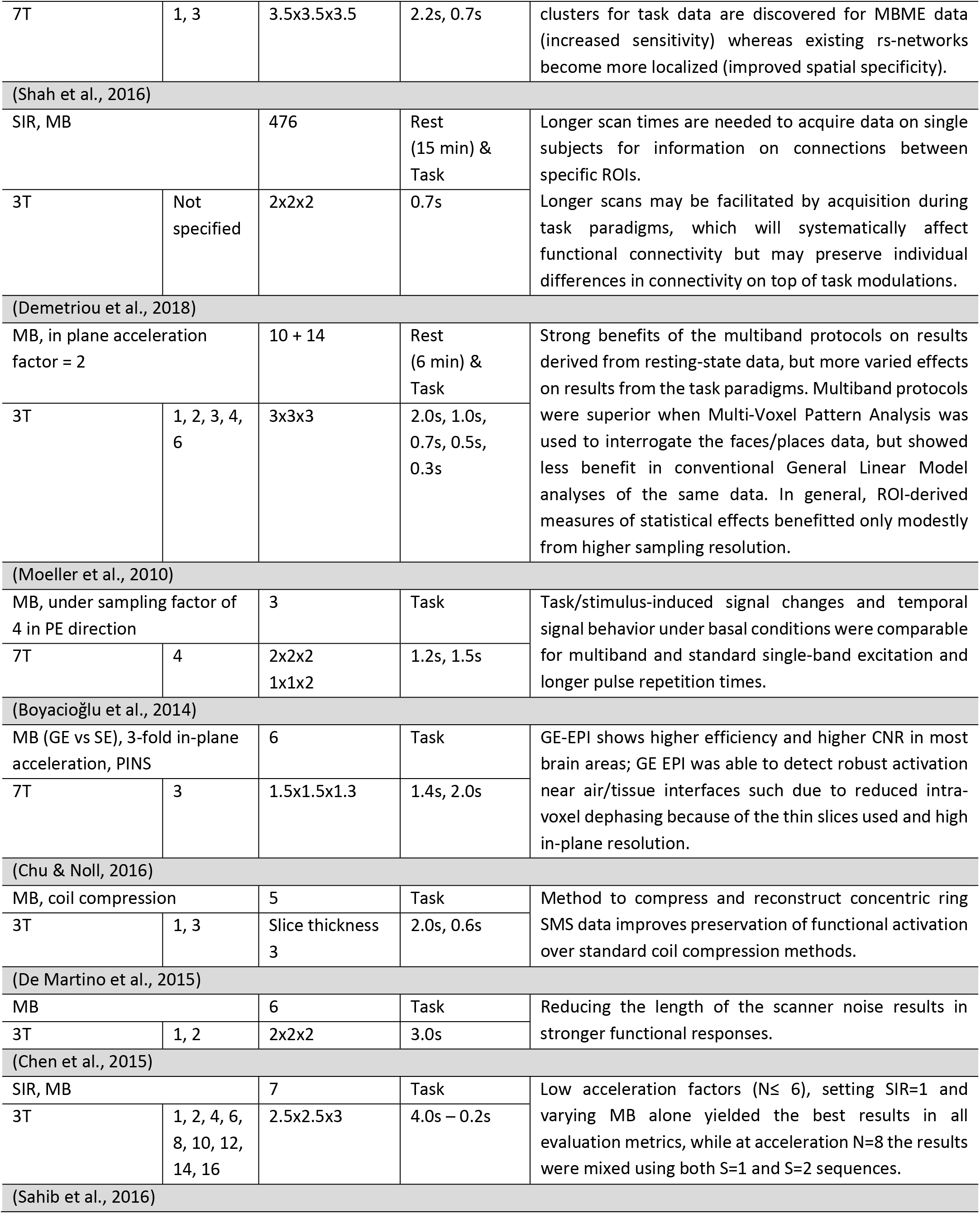
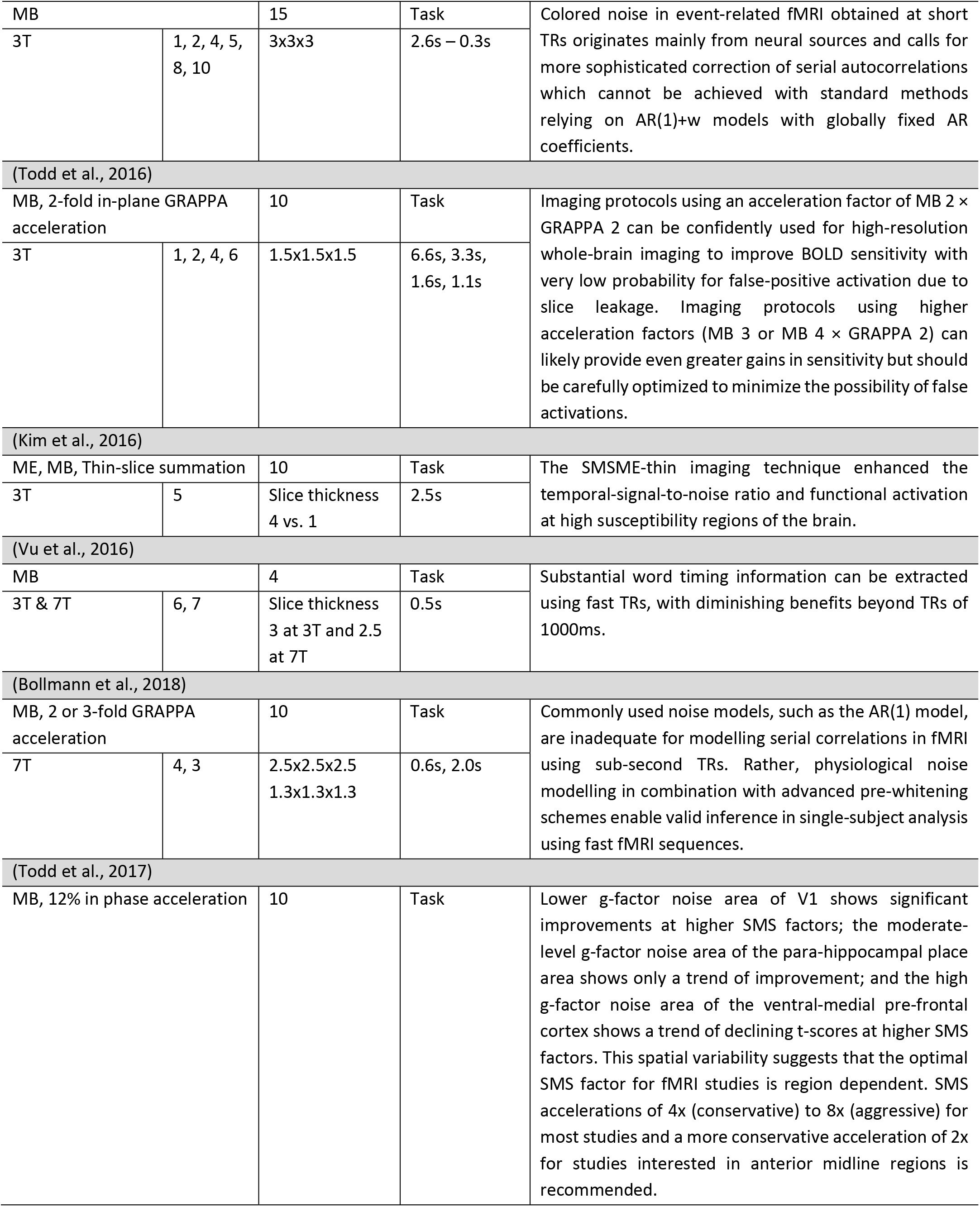
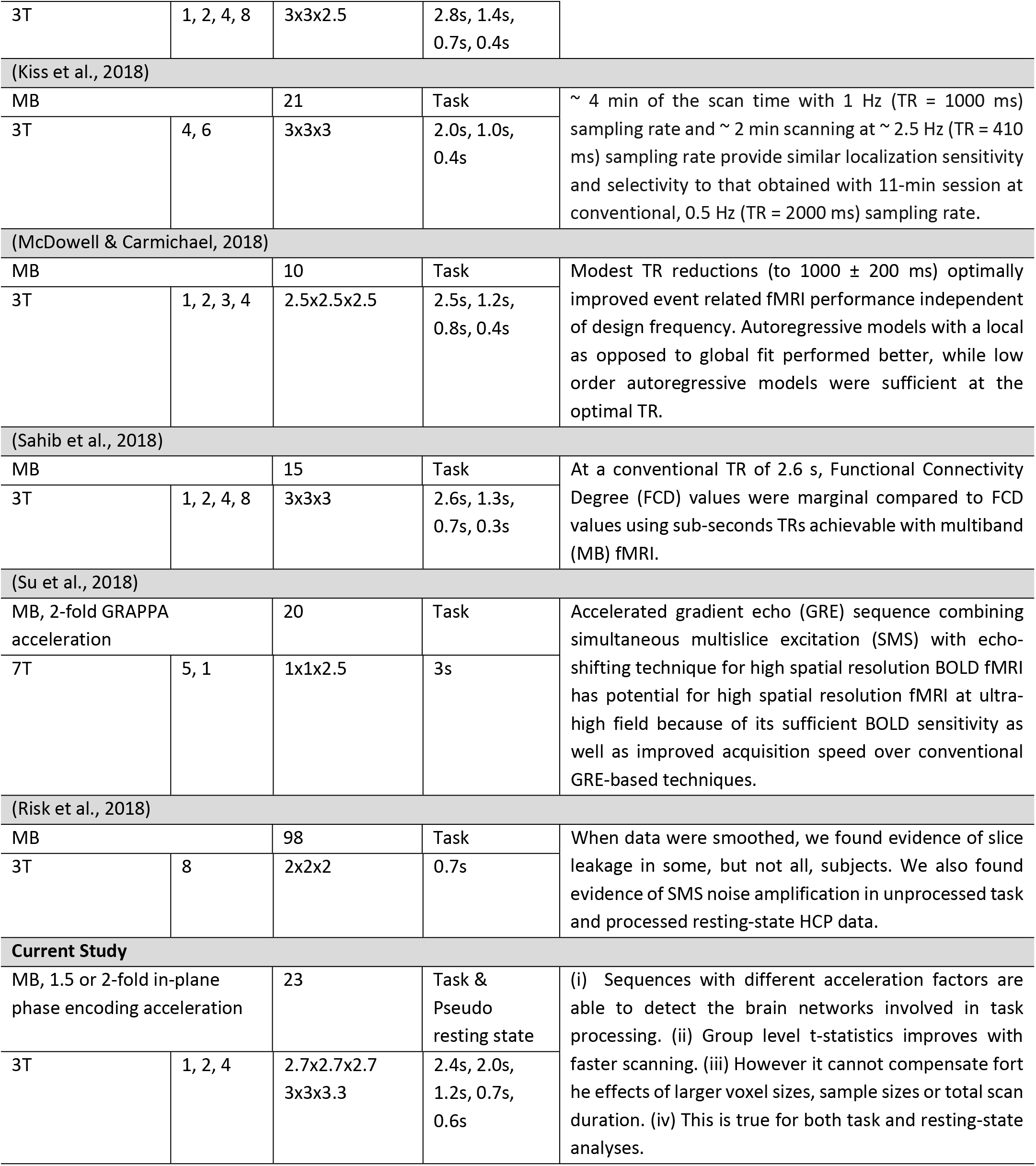
Summary of past studies exploring the benefits of MB acquisition and their main findings.

The main aim of the current study is to perform a detailed assessment of the effects of MB acceleration on random effects group level statistics for task-based fMRI, and identify the optimal acceleration for task-based fMRI. We therefore analysed the 20 studies that employed task based BOLD, in greater detail. Boyacioğlu et al., 2014 compared the performance of Gradient echo (GE) and Spin echo (SE) sequences in combination with MB factor of 3 and using a random effect analysis concluded that the performance of GE was superior in terms of the BOLD compared to SE. De Martino and colleagues, 2015, ingeniously used longer silent periods between two TRs to present their auditory stimuli and showed a higher BOLD contrast with MB2 compared to single band, concluding that reducing the length of the scanner noise results in stronger auditory responses. Similarly, other studies that used MB accelerated sequences, but did not directly compare their performances for task related statistics were not analysed further (Shah et al., 2016; Vu et al., 2016). We identified three other studies that directly compared MB sequences for noise amplification, slice leakage and serial autocorrelation matrices. These studies unequivocally showed that the noise is amplified as MB factor increases, leakage might occur at very high acceleration (>MB8) and that the conventional auto-correlation models might not be sufficient for data acquired with faster TRs (Bollmann, Puckett, Cunnington, & Barth, 2018; Boubela, Kalcher, Nasel, & Moser, 2014; Risk, Kociuba, & Rowe, 2018). Since group-level task-based BOLD was not the focus of these studies, we did not analyse them further. This left us with 13 studies which directly tested MB acquisition compared to the conventional single band acquisition or standard datasets to assess the benefits of higher sampling rate with small TRs on task-based fMRI. Collectively, these studies showed that using MB acceleration of 2 to 3 times might yield comparable or better statistics for task-based BOLD (Table 1). While these studies are valuable in enhancing our understanding of the effects of MB, there are several criteria, typical of modern cognitive neuroscience applications, not fulfilled in these studies and therefore warrants further investigation.

Up until now, the effects of MB acceleration have mostly been studied in basic functional designs using finger tapping and visual checkerboard (Chen et al., 2015; Chu & Noll, 2016; McDowell & Carmichael, 2018; Moeller et al., 2010; Sahib et al., 2018, 2016; Su et al., 2018; Todd et al., 2016). These tasks massively activate motor and visual cortices with strong contrasts between ON and OFF conditions. Since these are predominantly positioned in the cortex of the brain and are close to the MR receiver coils, higher acceleration factors are easily achieved. In contrast, modern cognitive neuroscience tasks have less statistical power and the networks are distributed beyond the cortex and throughout the brain, imposing a higher challenge in successfully accelerating reconstruction without impairing the signal. We thus feel that findings from MB-studies with basic sensory stimuli cannot be directly generalized to cognitive neuroscience applications. Moreover, while testing complex cognitive neuroscience applications, larger sample sizes are needed to reliably detect the BOLD as compared to basic stimuli where a sample size of ~15 is enough for a reliable activation. We therefore included a sample of 23 subjects to robustly assess MB-acceleration for complex stimuli.

Furthermore, the majority of these 13 studies that tested the effects of higher sampling rate only report summary statistics such as mean t- or top 10% t-values within selected ROIs from subject level analysis (Chen et al., 2015; Chu & Noll, 2016; Kiss, Hermann, Vidnyánszky, & Gál, 2018; McDowell & Carmichael, 2018; Sahib et al., 2018, 2016; Todd et al., 2017, 2016). Some studies additionally presented activation maps from representative subjects for visual comparisons (Moeller et al., 2010; Su et al., 2018), but only three studies presented voxel-wise, random-effects group-level, task-related BOLD statistics followed by a visual assessment of the performance of fast scanning.

In the current study, we add to our understanding of the effect of accelerated image acquisition and therefore higher sampling on task related BOLD statistics by addressing these gaps. We took advantage of a paradigm we have often used in our laboratory (Arnstein, Cui, Keysers, Maurits, & Gazzola, 2011; Gazzola & Keysers, 2009; Gazzola, Rizzolatti, Wicker, & Keysers, 2007; Valchev, Gazzola, Avenanti, & Keysers, 2016) and thus have reference data to systematically test the effect of MB acquisition. This paradigm aims to identify brain regions involved in the action observation network (Gazzola & Keysers, 2009) and involves observing **c**omplex goal directed **a**ctions (CA) and contrasting that activation against **c**omplex **c**ontrol stimuli (CC) that include the same objects and the same hand, but without the hand manipulating the objects. This contrast is motivated by the fact that viewing objects and hands in motion would already activate a very broad network including early visual areas, attentional brain circuits, spatial representations in addition to regions specifically representing how the hand manipulates the object. To focus on the latter, we thus subtract stimuli that include the same basic visual elements and their spatial distribution and also include movement, but without the actual hand-object interaction (Gazzola & Keysers, 2009). This contrast is known to identify a broad network of brain regions (often referred to as the action observation network) encoding hand-object interactions in all participants via the CA-CC contrast including premotor, somatosensory, insula, inferior parietal, visual, and cerebellar regions (Gazzola & Keysers, 2009). This allowed us to assess the benefits of MB across a wide range of brain regions and their functional connectivity. Unlike most of the previous studies, we present voxel-wise *group*-level analysis with detailed post-hoc analyses to quantify the differences in the data acquired with different acceleration using a sample size representative of modern task-based designs (n=23 subjects).

Based on the previous literature we hypothesized that the use of MB acquisition may show comparable if not higher t-values at a group level in response to task stimuli. Moreover, we performed a dual-regression analysis with the pseudo resting state data obtained by regressing out task-correlated activity to approximate the findings of the previous studies that showed an improvement of resting state statistics with MB acceleration. We hypothesized that on a group level, MB would outperform single-band acquisition in terms of the voxel-wise t-values.

## 2 Materials and methods

### 2.1 Subjects

Twenty-four (13 males, 11 females) right-handed (self-reported) healthy volunteers (M_age_ = 25.5 years, SD_age_ = 3.6, Range = 21 to 33) with no contraindications for MRI, and normal or corrected-to-normal vision participated in the study. After the screening procedure, subjects were familiarized with the MRI environment and were verbally informed about the observation task that they will watch during the session. The research was approved by the Amsterdam Medical Centre Ethics Committee Review Board, and the volunteers provided written consent at the start of the study. In what follows, conditions will be shorthanded as 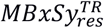, where TR is the acquisition time of one volume, res the resolution in mm, and x and y, the multiband and SENSE factors respectively. One subject (all sessions) was removed from the analysis due to excessive head motion and the 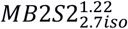 session of another subject was removed due to technical error during the acquisition, resulting in a final sample size of N=23 for 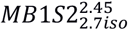, 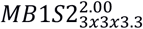, 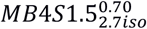, 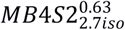 and of N=22 subjects for 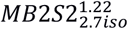 session.

### 2.2 Task and experimental procedure

Subjects were shown movie stimuli previously used in our lab to trigger robust activity in the action observation network (Valeria Gazzola & Keysers, 2009; Valchev et al., 2016). These movies show a human hand interacting with everyday objects. Examples of such interactions are shown in Figure 1 and listed in supplementary table 1 (see also Valchev et al., 2016). Two types of stimuli were used: Complex actions (CA) showed the hand interacting with the object in typical, goal directed actions. For example, the hand of the actor reached for a lighter placed on the table, grasped it, and lit a candle with it. Complex controls (CC) stimuli had the exact same setting as the CA but the actor’s hand did not interact with or manipulate the object on the table, instead, made aimless hand movements. A block was composed of three movies of the same category (CA or CC) and lasted 7s. Each fMRI session was composed of 13 blocks per stimulus category for a total of 26 blocks, presented in a randomized order. The inter-block-interval lasted between 8 – 12 s and consisted of a fixation cross on a gray and blue background similar to the stimuli background. Five of these sessions were presented to each subject, showing the same blocks but in different order, with each session acquired with a different acquisition scheme, varying in MB factor, in-plane SENSE (S) acceleration and/or spatial resolution (table 2). The order of acquisition was randomized between subjects. Importantly, the duration of the five sessions was similar (~8 min), but more functional volumes were acquired during sessions where the acquisition parameters lead to shorter TRs.

**Figure 1.**
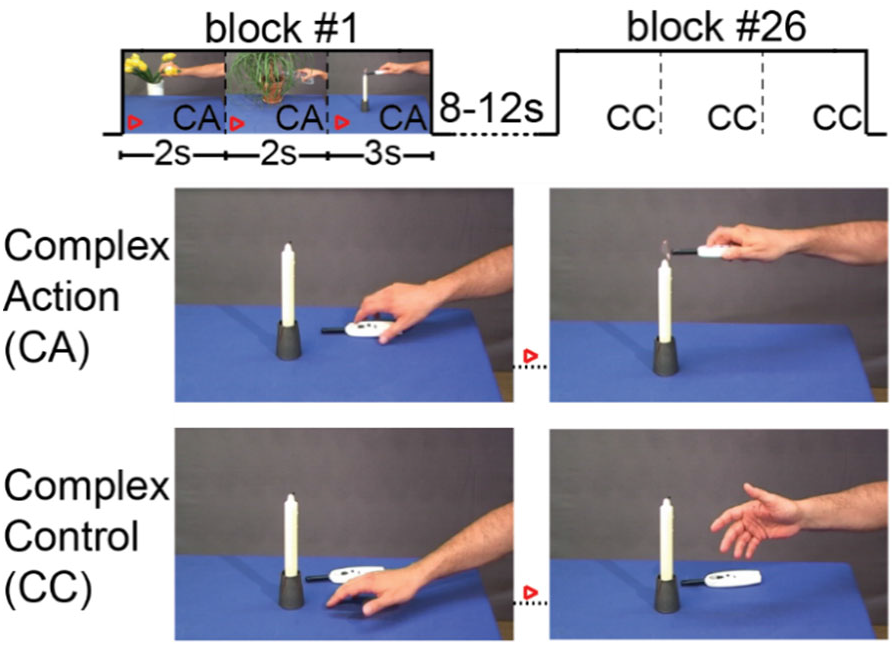
Subjects observed 26 blocks of video stimuli per session (x 5 acquisition sequences) showing one of the two conditions: complex action (CA) or complex control (CC). Thirteen bocks per condition were presented and each block comprised of three clips of the same condition. The inter block interval was randomized between 8 and 12 seconds. The blocks and conditions were randomized between the five acquisition sequences per subject and between subjects. The top row schematically illustrates the structure of each run; the bottom two rows illustrate, for a randomly chosen clip, how it differs in the two conditions: a goal directed action in CA and a random movement in CC.

**Table 2.**
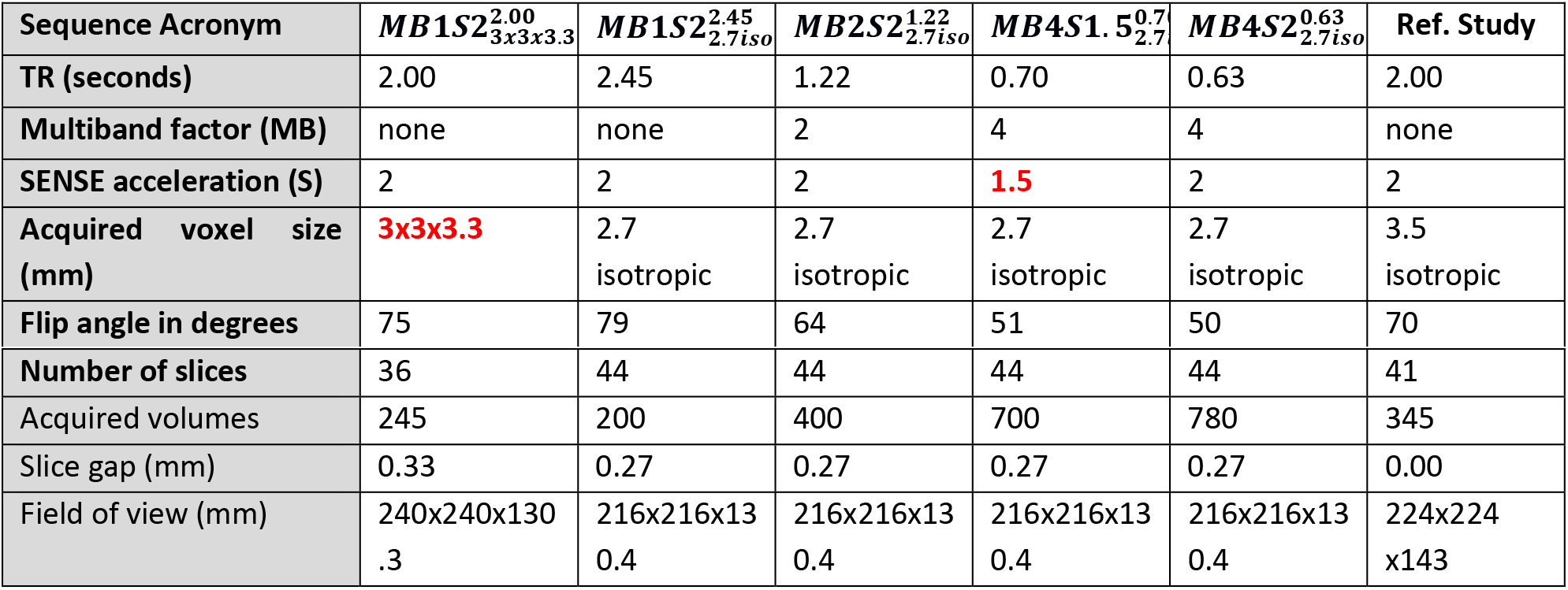
Overview of the scanning parameters used for the five acquisition sequences and the reference study. Note (in red) that the sequence 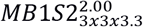 has a coarser spatial resolution compared to the other sequences. The sequence 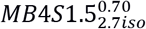 differs from 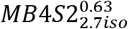 in the SENSE acceleration and not in MB factor. Reference study was collected with a different set of subjects and a different scanner. It is used to compare some results from this study.

### 2.3 Image acquisition

Data were acquired on a 3T Philips scanner, using a commercial version of Philips’ MB implementation (based on software release version R5.4). A 32-channel head coil was used. The total length of the scanning session per subject was around 50 minutes and included the functional and anatomical scans. Functional data were acquired using five different acquisition sequences (Table2). These sequences were chosen based on the following considerations. First, we wanted to acquire one of the most typically used non-multiband sequences as a meter of comparison, and therefore acquired a sequence with a TR=2.00s and close to 3mm isotropic resolution 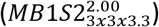. Because multiband is typically used with slightly smaller voxels, we acquired all other sequences at a resolution of 2.7mm isotropic, which is closer to the average gray matter thickness. At that resolution, we then measured an acquisition at MB1S2 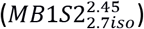, and increased MB to 2 and 4. At MB4, we additionally reduced SENSE acceleration to 1.5, to mitigate potentially higher noise amplification. The resulting sequences were 2.45s, 2.00s, 1.22s, 0.70s and 0.63s for the sequences 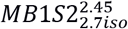, 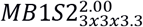, 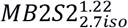, 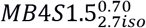 and 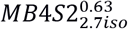 respectively (Table 2). Slices were acquired in ascending order in transverse direction. TE was kept constant for all sequences at 30ms. A default shift of ½ a FOV was used while acquiring with MB 2 and 4 to improve reconstruction quality. The flip angle was optimized to the Ernst angle of each TR. All functional images were reconstructed using the SENSE reconstruction algorithm (Setsompop et al., 2012) and this was done simultaneously with acquisition. Note that the sequence 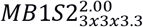 has a coarser spatial resolution compared to the other sequences, which would lead to differences that are not related to the total acceleration in acquisition. We therefore present the results of this sequence separately on figures rather than in the same line as the other sequences. In addition, the sequence 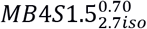 differs from 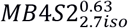 in the SENSE acceleration but not in MB factor. Any differences in these two sequences therefore will not reflect the effect of MB acceleration. A T1-weighted image was acquired at the end of the functional runs with a field of view of 240 x 256 x 250 mm and the voxel size was 1mm isotropic. Geometry factor (g-factor) maps (see supplementary analysis) that quantify the aliasing noise per voxel (Pruessmann, 1999) were obtained for sequences 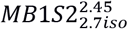, 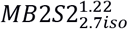, 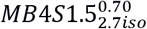. Here the g-factor maps were computed on-scanner using the vendor’s implementation and represent additional SNR penalty of multiband and in-plane SENSE acceleration. Note that the g-factor is computed from the coil sensitivity maps and the noise correlation matrix, both of which are measured in the preparation phase ahead of scanning (Pruessmann, 1999).

### 2.4 Reference study

As a reference for some of the analyses performed to test the effect of accelerated acquisition, we used a previously acquired dataset. Data from this previous study (Abdelgabar et al., 2018) consisted of fMRI data from an independent group of 31 subjects. Images included an anatomical scan and a functional run where the subjects viewed the same action observation paradigm used in this study (CA and CC blocks of 7s). In addition, they also saw thirteen 7s blocks of Static Control (SC) movies, which had the same elements as the CA and CC movies, but the hand lay still next to the objects without any movement. Data were acquired using a Philips Intera 3T scanner (University Medical Centre Groningen, University of Groningen. Groningen, The Netherlands), using a 32-channel coil. First, a high-resolution, structural image (170 slices; scan resolution = 256 x 256; field of view = 232 x 232 mm; voxel size: 1mm isotropic) was acquired. Functional images were acquired using an echo planar T2*-weighted gradient sequence (See Table 2 for additional parameters). The aim of choosing this independent dataset as a reference instead of the average data acquired in the current study was to avoid circularity, which may result in inflated statistical estimates, and to use a study with an even larger sample size to further approximate the true population activity that each multiband experiments aim to estimate. This data was acquired without any MB acceleration.

### 2.5 Preprocessing

Reconstructed whole brain functional data were preprocessed using SPM12 (Wellcome Trust Centre for Neuroimaging, UCL, UK) with Matlab version 8.4 (The MathWorks Inc., Natick, USA). Briefly, functional images were slice-time corrected and then realigned to the estimated average. Anatomical images were co-registered to the mean functional image, and segmented. The normalization parameters that were generated during segmentation were used to bring all the images to the MNI space. The resampled voxel size for the functional images was 2 × 2 × 2 mm and 1 x 1 x 1 for the anatomical scans. We adjusted the SPM12 bounding box settings to [−90 −126 −72; 90 90 108] in order to include the cerebellum completely, as the default settings of [−78 −112 −70; 78 76 85] may result in omission of some of the cerebellar voxels (Abdelgabar et al., 2018; Valeria Gazzola & Keysers, 2009). Smoothing of 6 mm FWHM Gaussian kernel was applied to the functional data. The reference dataset was preprocessed similarly.

### 2.6 Subject level General Linear Models (GLM)

#### GLM with task predictors

Subject level GLM included CA and CC as two separate task predictors with each predictor having 13 blocks of 7s. The reference dataset additionally included SC as a predictor with 13 blocks of 7s. Boxcar functions were convolved with canonical hemodynamic response function implemented in SPM. Regressors included the six motion parameters estimated during realignment, first five principal components of cerebrospinal fluid (CSF) and five principal components of white matter (WM) (total 16 regressors). The CSF and WM principal components were extracted from the normalized but unsmoothed functional data for each subject using the average WM and CSF segments of the subjects’ segmented and normalized anatomy, thresholded at 0.7 (arbitrary units) as masks (Behzadi, Restom, Liau, & Liu, 2007). We used the principal component analysis based regressors as they have been shown to yield better results for physiological de-noising as compared to regressors generated based on cardiac and respiratory traces recorded with the respiration belt and pulse oximeter (Kirilina, Lutti, Poser, Blankenburg, & Weiskopf, 2016). FAST autocorrelation algorithm implemented in SPM12 was used to correct for temporal autocorrelations in the data which has been suggested to be superior to the default AR(1) models specially for the smaller TRs (Bollmann et al., 2018; Todd et al., 2016). All the other parameters, including the high-pass filter (128 s), were left as the default in SPM12.

#### GLM for resting state

Previous studies with MB show that accelerating image acquisition may have benefits for detecting RSNs (Demetriou et al., 2018; Griffanti et al., 2014; Preibisch et al., 2015; Reynaud et al., 2017). To replicate these positive findings of MB acceleration on RSNs, in this study we created a pseudo resting state dataset by regressing out the BOLD associated with task predictors CA, CC and the 16 nuisance regressors. Briefly, a subject level GLM was performed on unsmoothed functional data with the GLM described above (i.e. two task predictors, CA and CC, and the 16 regressors). The residuals were saved which were devoid of the BOLD that was detected as correlated to our action observation task. These residuals were smoothed with 6mm FWHM Gaussian kernel and used in a spatial GLM in FSL (http://www.fmrib.ox.ac.uk/fsl/index.html) with 20 resting state networks (RSN20) as predictors (Smith et al., 2009). FSL was used as it has a well-implemented spatial GLM algorithm as a first step of the dual regression approach. The RSN20 maps classified by Smith et al consist of 10 primary cortical networks, 3 cortical networks with partial spatial/functional overlap with primary cortical network(s), 3 networks with spatial location corresponding to subcortical or deep cortical areas and 4 artefactual components. This spatial GLM step generated 20 time-courses per subject per acquisition sequence, one for each RSN. These time courses were then used as predictors in a subject level GLM along with the 16 nuisance regressors for the normalized and smoothed data. Note that this analysis is very similar to the dual regression analysis implemented in FSL. We decided to perform the second step in SPM to keep the default processing (e.g. auto-regression) similar to those used in the task-based GLM which was performed in SPM, so as to avoid effects of using different packages and to enable qualitative comparisons between task based and resting state analyses.

### 2.6 Group statistics

Random effects group t-tests were performed separately for each acquisition sequence using the parameter estimates from the subject level GLMs (CA-CC contrast or RSNs). The significance of all group analyses was evaluated at q_fdr_<0.05 at the voxel level and a cluster-size threshold of 50 voxels. We use FDR correction rather than FWE correction here to improve sensitivity (Lieberman & Cunningham, 2009; Lindquist & Mejia, 2015), but also provide histograms of t-values to enable scientists to appreciate the effect of MB at any threshold.

Basic signal and noise characteristics of different acquisition sequence, which include g-factor, tSNR and CNR are presented in supplementary materials (see Supplementary analysis). In the result section 3.1, we focus on the random effect group level task based BOLD outcomes for different acquisition sequences. We look at:

a. Voxel-wise t-statistics of individual sequences to see which sequences perform the best in terms of t-statistics. Next, using Receiver Operating Characteristics (ROC) on the resulting voxel-by-voxel t-values we analyze how similar/different the activation maps are between different acquisition sequences and the reference study. The ROC analysis allows us to look at the hit rate and false alarm rate in a non-threshold dependent manner. The intent of using the ROC measures is not to find a “winner sequence” but to see concordance between sequences. If a sequence would show poor concordance with all other sequences, it might warrant particular prudence. However, if all sequences show high concordance, this will point to the fact that although the t-statistics might differ between sequences, the sequence that one chooses would not greatly alter functional interpretation about the brain networks involved in the cognition of interest.
b. We then label the activated brain regions using the AAL atlas (Automated Anatomical Labeling, Tzourio-Mazoyer et al., 2002), and explore the similarity of conclusions at the brain-region level using an ROC analysis (i.e. which AAL region are detected as active and which not).
c. To explain the small differences in the outcomes of different acquisition sequences we performed within and between subject variance analyses to see if there are systematic differences in the variance patterns between different sequences as that would directly impact the observed t-statistics.
d. Previous studies with resting state show that MB acceleration may enable us to shorten total acquisition times (Smitha et al., 2018). To test this theory for task-based studies, we truncated the total acquired data and looked if it would be possible to reduce the number of task trials/total acquisition time with less loss in accuracy at higher MB acceleration.

Section 3.2 reports the group statistics of the pseudo resting state data as replication for the widely reported positive effects of MB in the resting state literature.

## 3 Results

### 3.1 Random effect group level analysis

#### (a) Voxel wise analyses

Figure 2A-B presents the random-effect group activation t-maps for each sequence separately. As a validation, we also present the group maps of the reference study (Figure 2C). Visual comparison of the group voxel-wise results look very similar across sequences, and consistent with what has been reported in Gazzola & Keysers (2009). Figure 2D presents the histograms of the significant t-values for the sequences 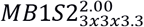, 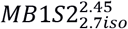, 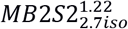, 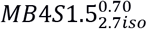 and 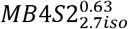. Both sequences with MB4 have higher t-values than sequences with MB 1 and 2 with same spatial resolution of 2.7mm isotropic.

**Figure 2.**
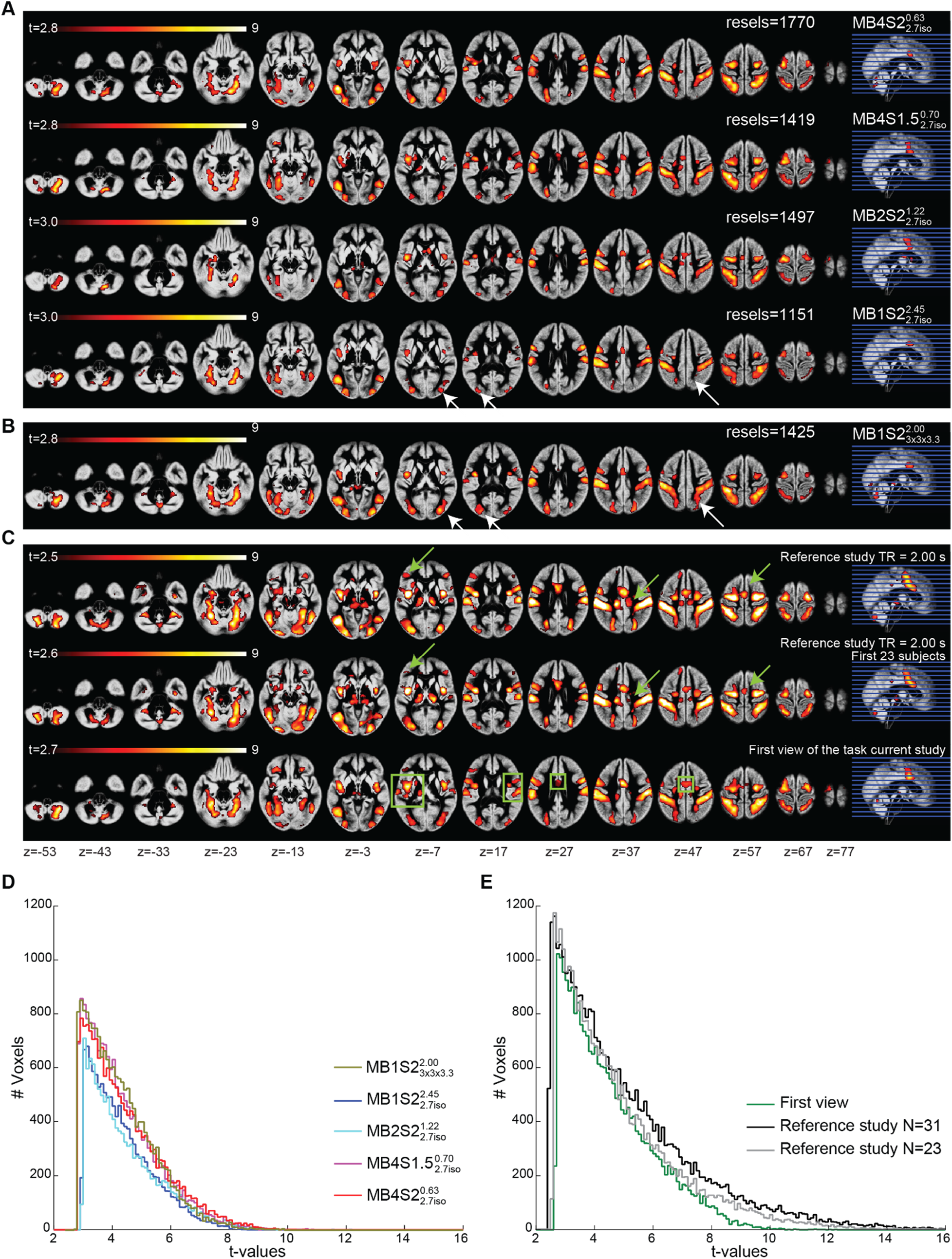
All maps overlaid on the mean grey matter segment of the group. q_fdr_<0.05, cluster threshold 50 voxels. **(A)** Group maps showing the task correlated activity detected using the GLM predictors for the acquisition sequences 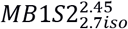, 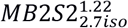, 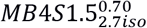 and 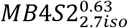. **(B)** Group maps for the acquisition sequence 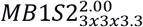. White arrows represent the effect of voxel size on the BOLD outcomes. **(C)** Group maps from the reference study using the same task (N=31 subjects), maps from the reference study with a smaller sample of N=23 subjects and from the current study looking at the first view of the task. Green arrows show how results change with the number of subjects. Green boxes represent the clusters that become bigger or more significant if we only consider the first view. **(D)** Histogram of the group t values for the CA-CC contrast in figure 2 A and B. **(E)** Histogram of the group t values for the CA-CC contrast in figure 2C.

One notable difference in the visual inspection of the maps is that all of the sequences tested here activate fewer voxels than the independent study when visualized at the same threshold (q<0.05). Figure 2E shows that the independent study (black line) has much higher t-values than the two best sequences (pink and red lines in Figure 2D) of the current study. This could potentially be due to three reasons: i) the independent study was conducted on 31 participants instead of the 23 participants included here, affording it more statistical power. ii) While in the reference study subjects viewed the stimuli only once, they saw them 5 times in our within-subject MB comparison, potentially leading to some form of repetition suppression. iii) The acquired voxel size in the reference study (3.5 mm isotropic) was larger than the resolution used here (2.7mm isotropic or 3×3×3.3mm). We therefore compared two more GLMs, one including just the first 23 subjects of the reference study and one with just the first view of the stimuli in the current study (Figure 2C). Note that the “first view” GLM consist of data coming from different sequences. To assess the impact of voxel size, we compared 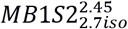 and 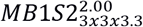, as these differ only in the acquired voxel sizes.

Visual inspection of the maps confirmed the intuition that the sample size (N=31 vs. N=23) has an impact on the group BOLD levels, as although the networks look similar, the size of the clusters appears a bit smaller when fewer participants are included (see green arrows in Figure 2C, and grey and black lines in Figure 2E). Next, we considered the first view in the current study regardless of the acceleration with which it was acquired. Some clusters start to appear as significant (green boxes in Figure 2C compared to maps in Figure 2A-B), which were either not present or were smaller, when group analyses were done per sequence and therefore contained first to fifth view of the task. Figure 2E shows that reference study (N=23) vs. the first view in this study are very similar with slightly higher t-values for the reference study. Finally, the impact of higher voxel size can be seen in sequences 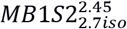 and 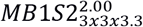, which differ in the voxel size but not in the acceleration: white arrows in Figure 2A-B show the areas with better BOLD for sequence with 3 × 3 × 3.3 mm^3^ voxels compared to sequence with 2.7 mm^3^ voxels (Figure 2D-E). Although it has been argued that temporal SNR no longer depends on voxel volume beyond a voxel-size of 1.5mm isotropic while using 32 channel coils (Triantafyllou, Polimeni, & Wald, 2011), our results show that going from 2.7mm^3^ isotropic to 3 × 3 × 3.3 mm^3^ improves the tSNR (Supplementary analysis 2) as well as the t-values for task based BOLD (Figure 2D). Taken together the findings of the random effect analysis suggest that in otherwise identical scanning parameters, MB4 shows higher t-values than lower acceleration with MB2 or no MB. Larger voxel sizes, larger group sizes and/or novelty of the task seems to show higher t-values. In our sample, MB1 sequence with larger voxels show t-values most comparable to MB4 sequences in terms of t-values, suggesting that MB4 can be used to obtain finer spatial resolution while keeping the t-sensitivity similar to a non-accelerated sequence with higher voxel sizes.

Since the GLM measures reported above are very threshold dependent, we used ROC measures to compare the conclusions one would reach from different acceleration factors. Table 3 presents signal detection metrics (ROC, hit rate and false alarm rate) comparing the outcomes of these GLMs. The area under the ROC curve, which is less threshold dependent, is very high in all pairwise comparisons. It ranges from 85% to 90% when comparing the sequences against the reference study, and from 94% to 98% when comparing different sequences acquired here. The hit-rate was low when assessing how many of the voxels in the reference study were activated in each sequence (40% on average), and there was no systematic trend of this hit rate increasing or decreasing with the total acceleration. The hit rate was much higher when examining how much of the voxels from the current sequences were activated in the highly powered reference study (83-90%), and when comparing different accelerated sequences with each other (72% on average). The false alarm rate was consistently low, with on average only 2% of the voxels not activated in the reference study activated at any sequences, and only 3% of voxels not activated in one sequence activated in another. The sub-sample (N=23) of the reference study showed high correspondence with the full sample (99%), had a high hit rate (81%) and a small false alarm rate (1%). The hit rate for the sequences against the subsample of the reference study was slightly higher (average 45%). The hit rate was also higher when only the first view (vs. the reference study) was considered. These matrices suggest that on average the maps from different acquisition sequences have a high concordance.

**Table 3.**
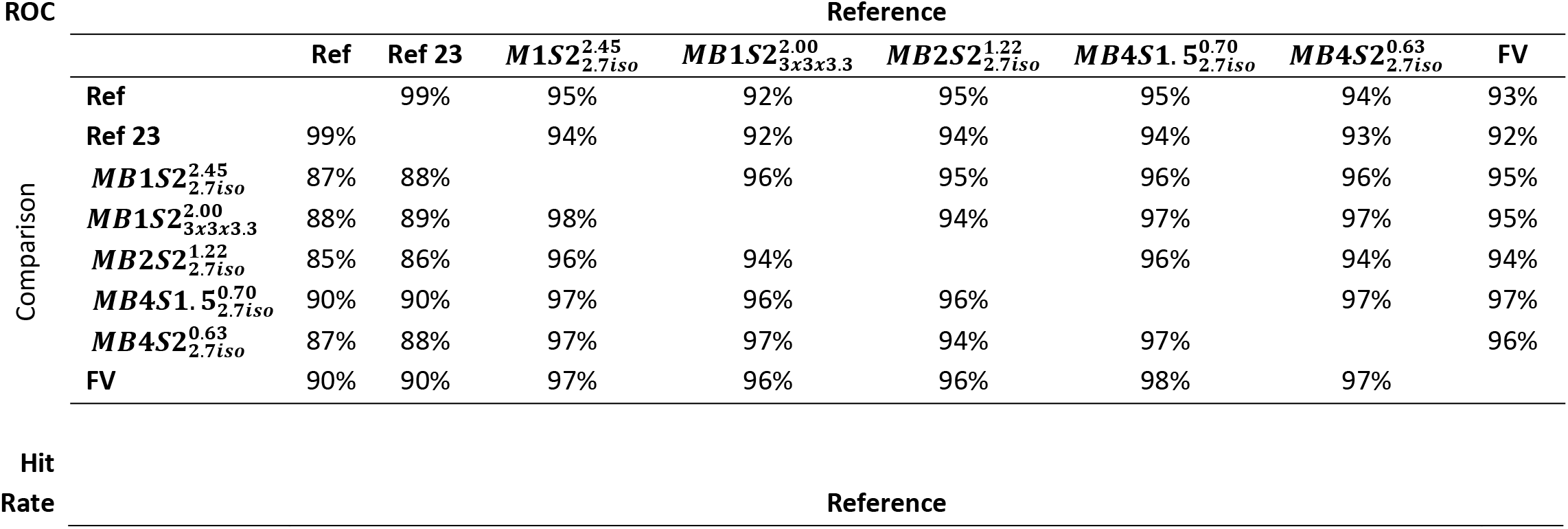
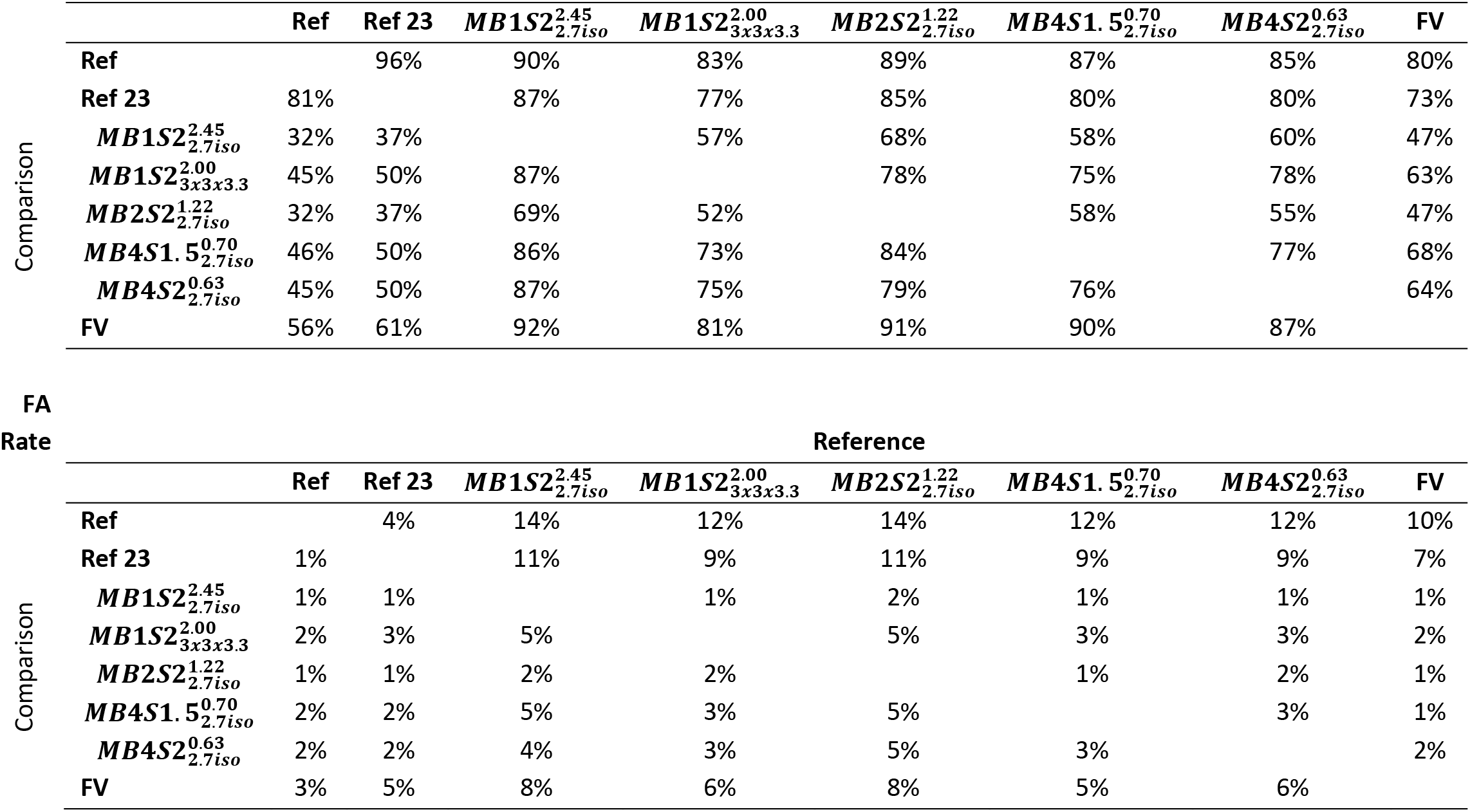
ROC measures for assessing similarity/differences between the t-maps resulting from the group ANOVA of the different sequences. Ref = reference study. Ref 23 = reference study with first 23 subjects. FV = first view in the current study.

#### (b) Region wise analyses

Next, we labelled the brain regions using AAL (thresholded at q_fdr_<0.05) to explore whether different sequences would lead to different conclusions about the brain network recruited by our task (Table 4). What is apparent from the table is that the core regions typically conceived of as part of the action observation network are found to be activated whatever parameters one uses (e.g. Postcentral, Precentral, Rolandic, and Supramarginal Gyrus). Such perfect agreement on the fact that a region is activated was true for 14 out of 62 brain regions in at least one hemisphere. There is also broad agreement on the fact that many regions are not activated (i.e. activated to less than 5%) in any of the sequences. This was true for 39 of the 62 brain regions in at least one hemisphere. In total, for 53 out of the tested 62 brain regions, scientists would thus arrive at the same conclusion whatever MB choice they would use. There was however a number of regions on which the different parameter choices did show substantial disagreement (highlighted in blue). It is notable, that regions of disagreement are often subcortical, including the cerebellum and basal ganglia.

**Table 4.** AAL labelling of the brain regions activated for the CA-CC contrast at q<0.05 level. Red colours are used to colour-code the level of activation in a region in terms of percentage activated. Blue cells are the brain region where there is a disagreement between different sequences, with some sequences showing more and some less than 5% activation (in both hemispheres). Note that for the vermis we show the joint activity of right and left hemisphere on the left side of the table. Values are % brain area active. Minimum red intensity is 5 and maximum is 50.

| Brain region | Left hemisphere |  |  |  |  | Right hemisphere |  |  |  |  |
| --- | --- | --- | --- | --- | --- | --- | --- | --- | --- | --- |
|  | *MB1S2<sup>2.00</sup><sub>3x3x3.3</sub>* | *MB1S2<sup>2.45</sup><sub>2.7iso</sub>* | *MB2S2<sup>1.22</sup><sub>2.7iso</sub>* | *MB4S1.5<sup>0.70</sup><sub>2.7iso</sub>* | *MB4S2<sup>0.63</sup><sub>2.7iso</sub>* | *MB1S2<sup>2.00</sup><sub>3x3x3.3</sub>* | *MB1S2<sup>2.45</sup><sub>2.7iso</sub>* | *MB2S2<sup>1.22</sup><sub>2.7iso</sub>* | *MB2S2<sup>1.22</sup><sub>2.7iso</sub>* | *MB4S2<sup>0.63</sup><sub>2.7iso</sub>* |
| Postcentral | 30 | 26 | 33 | 32 | 30 | 33 | 32 | 33 | 39 | 39 |
| Parietal Sup | 45 | 29 | 27 | 37 | 39 | 29 | 14 | 16 | 25 | 20 |
| Occipital Inf | 39 | 34 | 30 | 34 | 42 | 24 | 13 | 11 | 13 | 18 |
| Parietal Inf | 35 | 29 | 27 | 34 | 36 | 25 | 17 | 14 | 20 | 23 |
| SupraMarginal | 30 | 28 | 21 | 39 | 34 | 23 | 18 | 14 | 26 | 22 |
| Cerebelum 6 | 38 | 26 | 6 | 21 | 25 | 33 | 29 | 16 | 24 | 36 |
| Occipital Mid | 34 | 20 | 14 | 20 | 32 | 27 | 13 | 14 | 18 | 30 |
| Fusiform | 32 | 22 | 17 | 28 | 24 | 14 | 8 | 1 | 9 | 9 |
| Precentral | 17 | 20 | 26 | 27 | 25 | 5 | 7 | 8 | 9 | 7 |
| Cerebelum 8 | 5 | 3 | 0 | 8 | 11 | 28 | 23 | 16 | 23 | 25 |
| Rolandic Oper | 18 | 11 | 17 | 17 | 24 | 6 | 4 | 17 | 16 | 8 |
| Occipital Sup | 26 | 6 | 6 | 16 | 23 | 25 | 0 | 0 | 9 | 11 |
| Insula | 19 | 16 | 12 | 26 | 19 | 5 | 4 | 6 | 4 | 4 |
| Frontal Inf Oper | 8 | 7 | 7 | 16 | 15 | 12 | 13 | 6 | 15 | 13 |
| Vermis 8 | 28 | 0 | 3 | 5 | 18 |  |  |  |  |  |
| Vermis 7 | 32 | 0 | 0 | 0 | 11 |  |  |  |  |  |
| Temporal Inf | 9 | 6 | 5 | 8 | 7 | 10 | 7 | 4 | 9 | 9 |
| Cingulum Mid | 5 | 6 | 13 | 14 | 8 | 4 | 2 | 4 | 5 | 2 |
| Cerebelum 4 5 | 5 | 3 | 4 | 6 | 4 | 9 | 6 | 3 | 9 | 10 |
| Cerebelum 7b | 2 | 1 | 9 | 2 | 3 | 8 | 8 | 0 | 12 | 11 |
| Vermis 6 | 8 | 0 | 0 | 0 | 16 |  |  |  |  |  |
| Thalamus | 1 | 0 | 4 | 14 | 0 | 0 | 0 | 7 | 0 | 0 |
| Frontal Sup | 6 | 5 | 7 | 12 | 9 | 5 | 5 | 7 | 8 | 7 |
| Lingual | 13 | 2 | 1 | 8 | 6 | 2 | 0 | 1 | 2 | 2 |
| Cerebelum 9 | 0 | 0 | 0 | 0 | 0 | 13 | 16 | 0 | 0 | 8 |
| Putamen | 0 | 5 | 0 | 21 | 4 | 0 | 1 | 0 | 0 | 5 |
| Frontal Sup Orb | 0 | 0 | 0 | 0 | 0 | 0 | 0 | 0 | 0 | 0 |
| ParaHippocampal | 3 | 2 | 6 | 3 | 3 | 2 | 1 | 0 | 2 | 1 |
| Supp Motor Area | 0 | 0 | 8 | 8 | 0 | 0 | 0 | 3 | 0 | 0 |
| Pallidum | 0 | 4 | 0 | 11 | 4 | 0 | 0 | 0 | 0 | 0 |
| Hippocampus | 2 | 0 | 6 | 0 | 1 | 0 | 0 | 0 | 0 | 0 |
| Caudate | 0 | 0 | 3 | 0 | 0 | 0 | 0 | 6 | 0 | 0 |
| Heschl | 1 | 0 | 5 | 0 | 3 | 0 | 0 | 0 | 0 | 0 |
| Temporal Mid | 5 | 4 | 2 | 4 | 4 | 5 | 4 | 3 | 4 | 4 |
| Cerebelum Crus1 | 2 | 1 | 0 | 0 | 1 | 0 | 1 | 0 | 0 | 3 |
| Precuneus | 3 | 0 | 1 | 4 | 4 | 1 | 0 | 0 | 1 | 1 |
| Temporal Sup | 2 | 2 | 2 | 2 | 4 | 0 | 0 | 1 | 1 | 0 |
| Amygdala | 0 | 4 | 4 | 0 | 0 | 0 | 0 | 0 | 0 | 0 |
| Cingulum Ant | 1 | 0 | 1 | 3 | 2 | 0 | 0 | 0 | 1 | 0 |
| Vermis 10 | 4 | 0 | 0 | 0 | 0 |  |  |  |  |  |
| Frontal Mid Orb | 0 | 0 | 0 | 1 | 0 | 0 | 0 | 0 | 0 | 0 |
| Calcarine | 2 | 0 | 0 | 0 | 0 | 1 | 0 | 0 | 0 | 1 |
| Frontal Inf Orb | 0 | 0 | 0 | 4 | 0 | 0 | 0 | 0 | 0 | 0 |
| Angular | 0 | 0 | 0 | 0 | 0 | 4 | 0 | 0 | 1 | 0 |
| Cerebellum Crus2 | 1 | 0 | 0 | 0 | 1 | 1 | 0 | 0 | 0 | 1 |
| Frontal Mid | 0 | 0 | 1 | 1 | 1 | 1 | 1 | 1 | 2 | 1 |
| Cuneus | 0 | 0 | 0 | 0 | 0 | 2 | 0 | 0 | 0 | 1 |
| Paracentralobule | 0 | 0 | 1 | 1 | 0 | 0 | 0 | 0 | 0 | 0 |
| Frontal Inf Tri | 0 | 0 | 0 | 0 | 0 | 0 | 0 | 0 | 0 | 0 |
| Cerebellum 10 | 0 | 0 | 0 | 0 | 0 | 1 | 0 | 0 | 0 | 0 |
| Vermis 1 2 | 0 | 0 | 0 | 0 | 0 |  |  |  |  |  |
| Vermis 3 | 0 | 0 | 0 | 0 | 0 |  |  |  |  |  |
| Vermis 4 5 | 0 | 0 | 0 | 0 | 0 |  |  |  |  |  |
| Vermis 9 | 0 | 0 | 0 | 0 | 0 |  |  |  |  |  |
| Cerebellum 3 | 0 | 0 | 0 | 0 | 0 | 0 | 0 | 0 | 0 | 0 |
| Cingulum Post | 0 | 0 | 0 | 0 | 0 | 0 | 0 | 0 | 0 | 0 |
| Frontal Med Orb | 0 | 0 | 0 | 0 | 0 | 0 | 0 | 0 | 0 | 0 |
| Frontal Sup Medial | 0 | 0 | 0 | 0 | 0 | 0 | 0 | 0 | 0 | 0 |
| Olfactory | 0 | 0 | 0 | 0 | 0 | 0 | 0 | 0 | 0 | 0 |
| Rectus | 0 | 0 | 0 | 0 | 0 | 0 | 0 | 0 | 0 | 0 |
| Temporal Pole Mid | 0 | 0 | 0 | 0 | 0 | 0 | 0 | 0 | 0 | 0 |
| Temporal Pole Sup | 0 | 0 | 0 | 0 | 0 | 0 | 0 | 0 | 0 | 0 |

While this provides a qualitative overview, to get a quantitative summary measure of how similar a conclusion would be drawn from such activation tables using different sequences, we examined ROC measures and correlation measured across the regions obtained using different sequences (Table 5). The ROC analysis revealed that the concordance has an area under the curve of at least 0.87 for the worst of the comparisons, and is very close to perfect (0.99) when comparing the highest acceleration 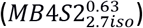 against the gold standard sequence without MB 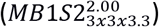 that had voxels with 50% larger volume. Correlation measures on un-thresholded tables (in terms of percent activated) was at least 0.77 and on average 0.88, further supporting how similar a conclusion would be reached. Binarizing the table in those regions activated (i.e. with at least 5% of the voxels in that region showing significant activation) or not, reduced the correlation to 0.74 on average.

**Table 5.** ROC and correlations between the levels of activation labelled using AAL atlas. No threshold in Yellow and with 5% threshold in green.

| ROC |  | Reference |  |  |  |  |
| --- | --- | --- | --- | --- | --- | --- |
| Comparison | | $MB1S2^{2.45}_{2.7iso}$ | $MB1S2^{2.00}_{3x3x3.3}$ | $MB2S2^{1.22}_{2.7iso}$ | $MB4S1.5^{0.70}_{2.7iso}$ | $MB4S2^{0.63}_{2.7iso}$ |
| | $MB1S2^{2.45}_{2.7iso}$ | | 0.93 | 0.89 | 0.94 | 0.93 |
| | $MB1S2^{2.00}_{3x3x3.3}$ | 0.95 | | 0.88 | 0.92 | 1.00 |
| | $MB2S2^{1.22}_{2.7iso}$ | 0.91 | 0.89 | | 0.90 | 0.87 |
| | $MB4S1.5^{0.70}_{2.7iso}$ | 0.96 | 0.92 | 0.92 | | 0.93 |
| | $MB4S2^{0.63}_{2.7iso}$ | 0.97 | 0.99 | 0.89 | 0.94 | |

Correlation
|  | *MB1S2*<sup>2.45</sup><sub>2.7iso</sub> | *MB1S2*<sup>2.00</sup><sub>3x3x3.3</sub> | *MB2S2*<sup>1.22</sup><sub>2.7iso</sub> | *MB4S1.5*<sup>0.70</sup><sub>2.7iso</sub> | *MB4S2*<sup>0.63</sup><sub>2.7iso</sub> |
| --- | --- | --- | --- | --- | --- |
| *MB1S2*<sup>2.45</sup><sub>2.7iso</sub> |  | 0.87 | 0.87 | 0.92 | 0.93 |
| *MB1S2*<sup>2.00</sup><sub>3x3x3.3</sub> | 0.77 |  | 0.77 | 0.84 | 0.94 |
| *MB2S2*<sup>1.22</sup><sub>2.7iso</sub> | 0.64 | 0.62 |  | 0.90 | 0.86 |
| *MB4S1.5*<sup>0.70</sup><sub>2.7iso</sub> | 0.78 | 0.77 | 0.63 |  | 0.92 |
| *MB4S2*<sup>0.63</sup><sub>2.7iso</sub> | 0.81 | 0.92 | 0.62 | 0.81 |  |

Taken together these findings suggest that on a group level, with the exception of a few regions, one would reach very similar conclusions about the cortical networks that are involved in action observation, regardless of which acquisition sequence is used, with the highest acceleration allowing a voxel-volume reduction of 50% without changing conclusions.

#### (c) Between and within subject variances

The measurements with MB acceleration are noisier due to the imperfect separation of the simultaneously acquired voxels (Golestani et al., 2017). However, it also allows acquisition of many more samples which are thought to increase the temporal SNR per time unit (see supplementary analysis 1, 2). As the noise levels have a direct impact on the t-value estimation, we looked at the between and within subject variances to see if the marginally higher t-values in sequences with MB4 can be explained by systematic differences in the variances. One might expect that MB acceleration has benefits on within subject variance by providing more samples, but not on between subject variance, because we did not measure more participants at higher MB. We therefore performed a variance analysis as explained in Kirilina, Lutti, Poser, Blankenburg, & Weiskopf, 2016. The maps of the intra-subject variance (σ^2^) were calculated using equation 3.1.

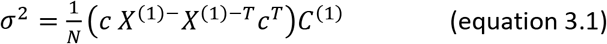

Here, N= number of subjects, c = contrast of interest (CA-CC), X^(1)-^ is the generalized inverse of the design matrix of the fixed effect analysis and C^(1)^ is the residual variance maps (ResMS) generated while estimating the fixed effect ANOVA. Superscripted T signifies the transposition of the matrix. Since the random effect variance maps are a combination of inter subject and intra subject variances, inter-subject variance maps (Σ^2^) were calculated by subtracting σ^2^ from the ResMS maps estimated during the random effect second level ANOVA. Five voxels with peak activities in the CA-CC contrast of the reference study (Figure 2C) were selected as centers of 6 mm radius spheres (ROI1: −50, −24, 34 area PF of the inferior parietal lobe; ROI2: 40, −32, 44 Area 2 of the primary somatosensory cortex; ROI3: 30, −12, 60 superior frontal gyrus; ROI4: −40, −6, 4 insula lobe; ROI5: 38, −4, 10 area OP3 of the secondary somatosensory cortex; See inset in figure 3 for the location of each ROI). Figure 3 presents the mean t-values for the fixed and random effect models, and within- and between-subject variances. As can be seen, some ROIs were dominated by within- (ROI4, 5) and some by between-subject (ROI1, 2) variance, but in all cases, the 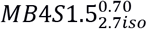 sequence outperforms the 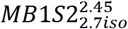 sequence in terms of t-values. Also, within subject variance was often but not always reduced in 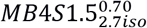 vs. 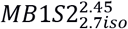 sequences. There was thus no systematic relationship between benefits of MB against the dominant source of variance.

**Figure 3.**
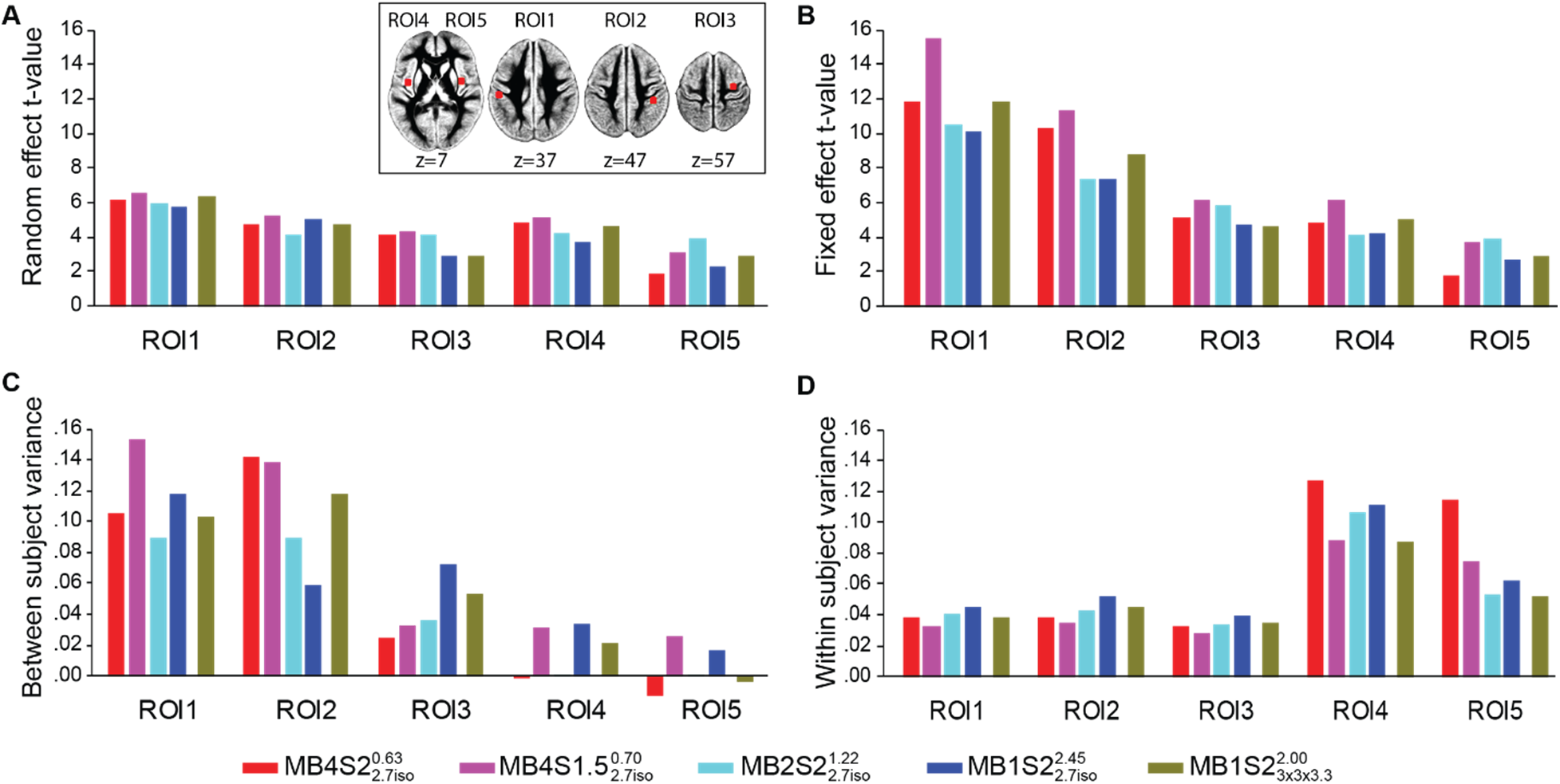
Mean parameter values for the CA-CC contrast from five ROIs (radius 6mm) centered on the first five voxels showing highest t values (at correction q_fdr_<0.05) for the CA-CC contrast in the reference study. Figure A and B show the mean t-values from the random effect model and the fixed effect model, respectively. Figure C and D show the between-subject variance and the within-subject variance in these ROIs, respectively. Inset shows the location of the five ROIs.

#### (d) GLM on first third of the data

With a specific total acquisition time, higher MB allows the acquisition of up to 4 times the number of volumes acquired with no MB. We thus explored whether the additional volumes per unit time at high MB can help when experimental time is limited. To test this, we repeated the subject level analysis using only the first third of each session for all five sequences. Group level t-maps were calculated using the CA-CC contrast image of each sequence separately and the resulting group t-map was correlated to the group t-map of the same contrast from the reference study. A similar correlation was also computed between the t-maps of the full GLM (see section 3.2) and the reference study.

As can be seen in the t-maps of the truncated dataset (one-third of the total scan) presented in Figure 4A, many nodes that are a part of the action observation network can already be detected (see Figure2A-C for comparison). Moreover, looking at the histogram of these images, the overall performance pattern is similar to the results of the full dataset: higher acceleration result in higher t-values (Figure 4B). 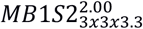 which have bigger voxel size also shows higher t-values than the sequence with same acceleration but smaller voxel size 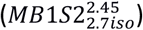. Looking at the correlation levels, as expected, none of the truncated dataset reaches the same level as the full dataset of the same sequence (Figure 4C), or as the full dataset of the MB1 sequence, suggesting that to obtain a reliable activation map, multiband is not a surrogate for total acquisition time.

**Figure 4.**
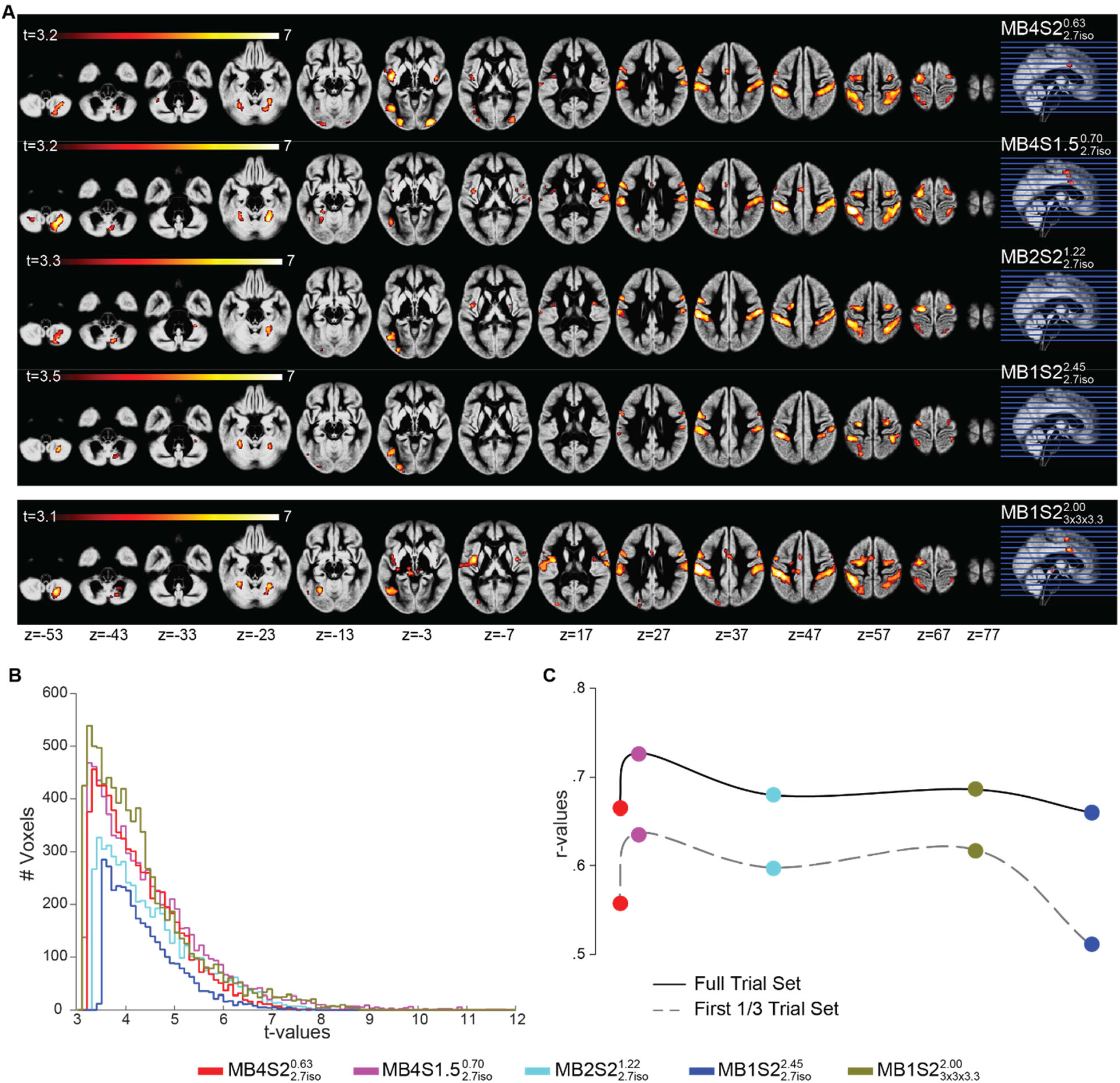
**(A)** Group maps showing the task correlated activity detected using the task GLM predictors, but using only the first one third of the total acquisition per sequence. Overlaid on the mean grey matter segment of the group. q_fdr_<0.05, cluster threshold 50 voxels. **(B)** Histogram of the t values for the CA-CC contrast for the one third of the total acquisition per sequence. **(C)** Correlation between t-maps of the CA-CC contrast per sequence and t-maps of the same contrast from the 31 subjects of the reference study.

### 3.2 GLM for Resting State Networks

Multiband has so far been mainly used and validated for resting state studies, where it has been suggested to improve the ability to detect RSNs. To explore whether we can replicate that finding in our data, we performed a pseudo resting state analysis with our data. The detailed description of the method can be found in section 2.6.

Figure 5 shows a representative network (rsn06 in Smith et al., 2009) separately for each sequence. Overall, at the threshold of q_fdr_<0.05, the network looks very similar for all the five sequences. Careful visual examination shows a small decrease in the cluster sizes as the TR increases (see white arrows and white boxes in Figure 5 for examples).

**Figure 5.**
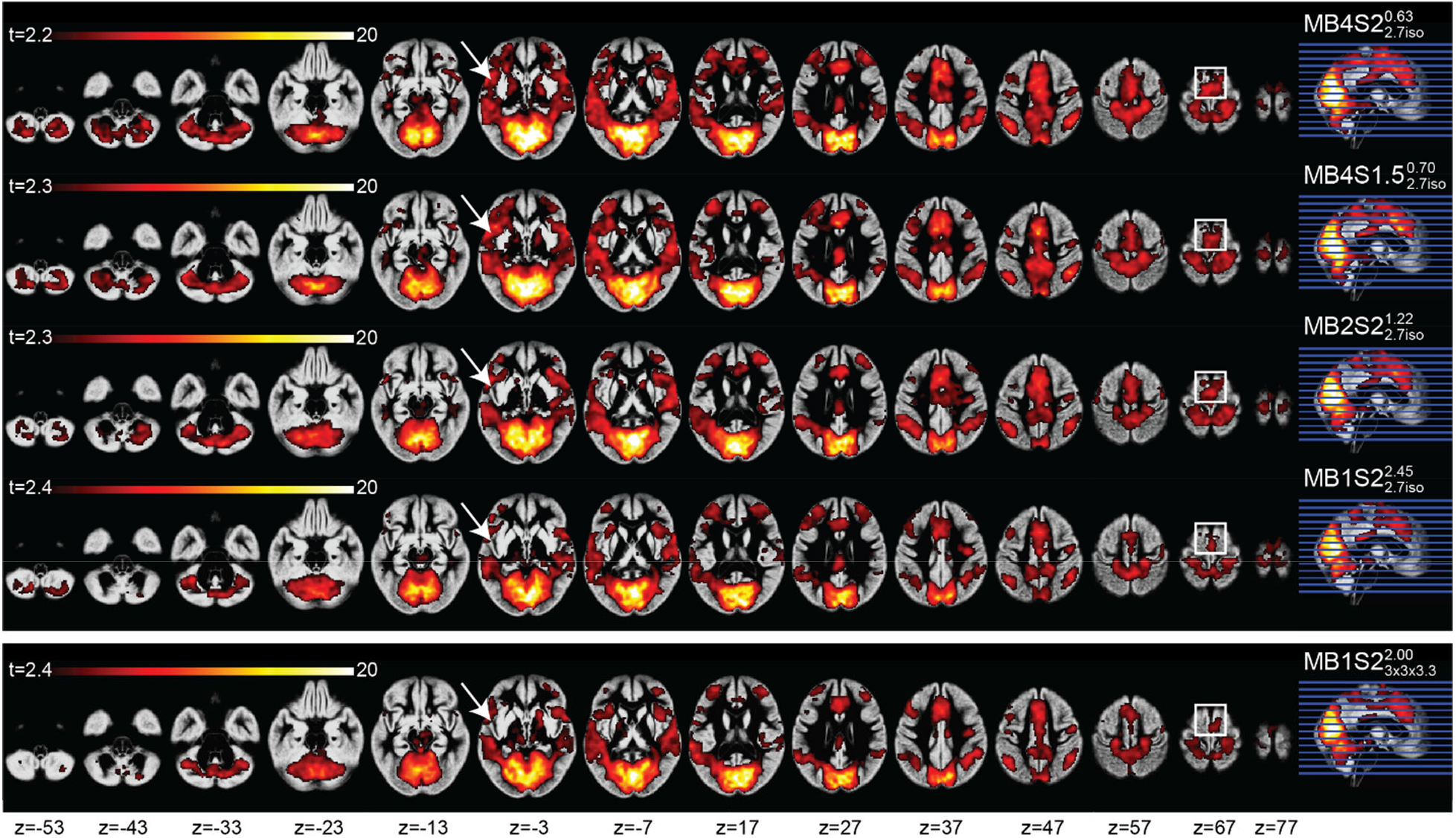
Group maps per acquired sequence showing a representative RSN: network number 6, as described in Smith et al. (2009). White arrows and rectangles evidence areas with visually different cluster extension across different sequences. Maps overlaid on the mean grey matter segment of the group, and thresholded at q_fdr_<0.05 with a minimum cluster size of 50 voxels.

To quantify this, we counted the total number of voxels that were significant at q_fdr_<0.05 for all 20 networks. Figure 6 present the average number (over all networks) of voxels as a function of acquisition sequence. An ANOVA was computed in SPSS with 20 components and 5 sequences and revealed a main effect of sequence (F(4,76)=28.82, p<0.001). Post-hoc pairwise comparisons show that the slowest acquisition 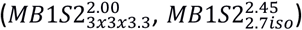 have fewer significant voxels than any of the MB accelerated sequences 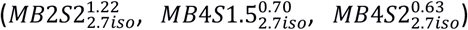. On average across all RSNs, sequence with 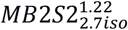 show a 15% increase in the number of significant voxels and sequence with 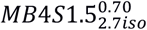 and 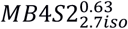 show 35% and 38% increase respectively, as compared to 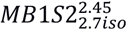, 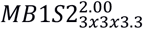 had the lowest number of voxels significant at q_fdr_<0.05.

**Figure 6.**
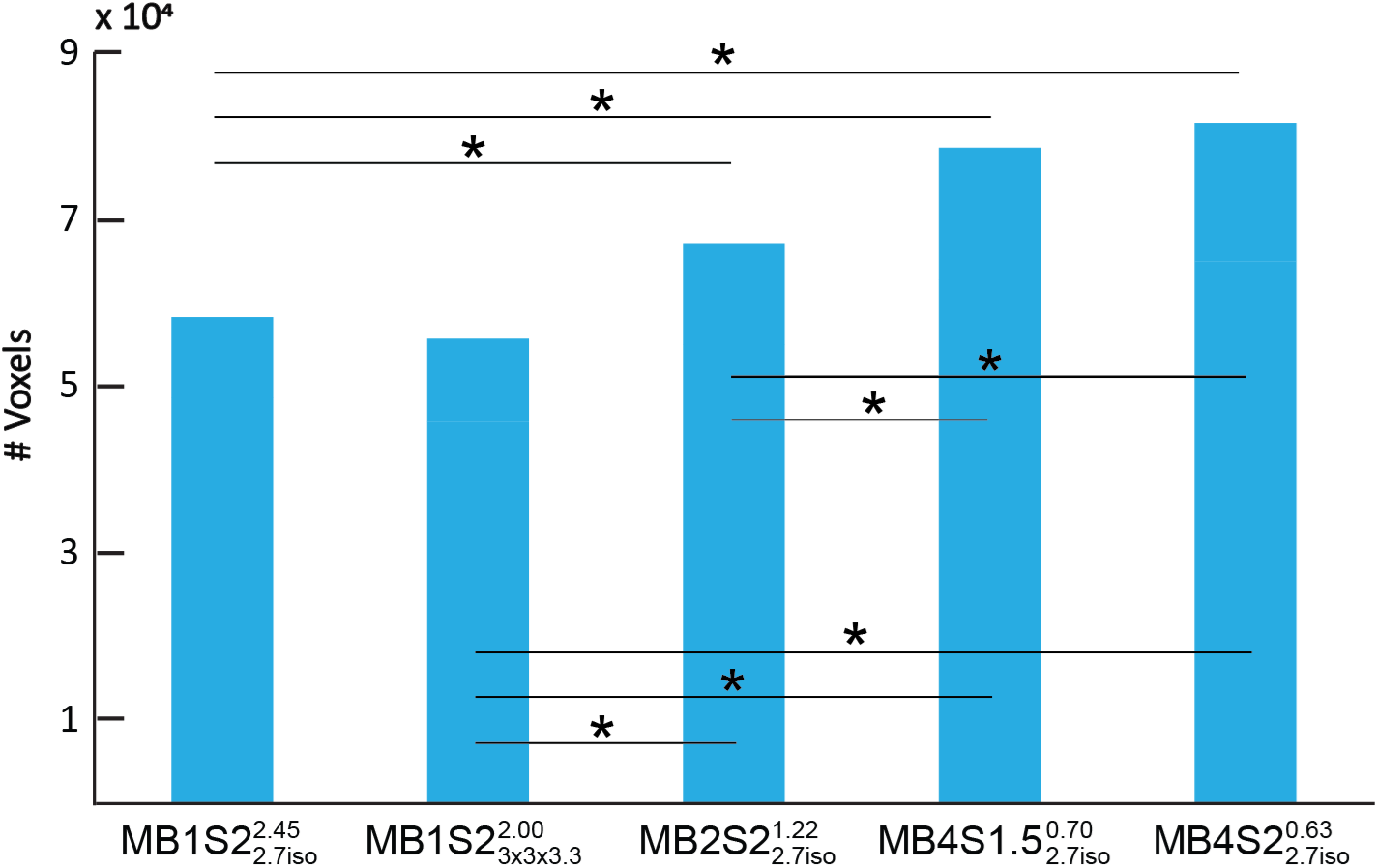
Total number of voxels that are significant at q_fdr_<0.05 for each sequence. (* p<0.001 for between sequence comparisons.)

To explore this phenomenon in a way that is less threshold dependent, and separately for each rsn, we plotted normalized cumulative histograms in Figure 7. Using q_fdr_<0.05 for each rsn for 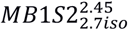, we counted the number of significant voxels, N_ref_. For any higher point along the x axis, we plot N(*t* ≥ x)/N_ref_. Accordingly, for 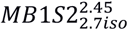, the first point along the x axis has a value of 1, which then decreases as a higher threshold is used while moving to the right. Values of other x-coordinates of sequences can then be directly understood as the number of voxels surviving that threshold relative to N_ref_. Plots framed in the orange outline show the component for which 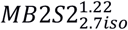 performs best, and plots framed in red are the ones where sequences with no MB show the highest number of supra-threshold voxels. All the others components showed that 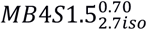 or 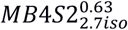 indeed identify more voxels across a wide range of thresholds. The two components (in red) which do not show an advantage of MB are classified by Smith et al., 2009 as belonging either to deep brain regions, or are artefactual components. However, since 17 out of 20 components show that using MB4 would yield higher t-values, this analysis recommends the use of MB 4 while studying RSNs at the group level if the aim is to be inclusive. One unexpected finding of this analysis was that the 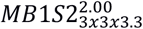, which was excellent for task based fMRI (performing better than 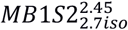 and similar to the 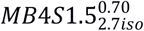 and 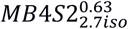), was one of the least sensitive sequence for the RSN. The potential reason for this curious finding is discussed in the discussion section.

**Figure 7.**
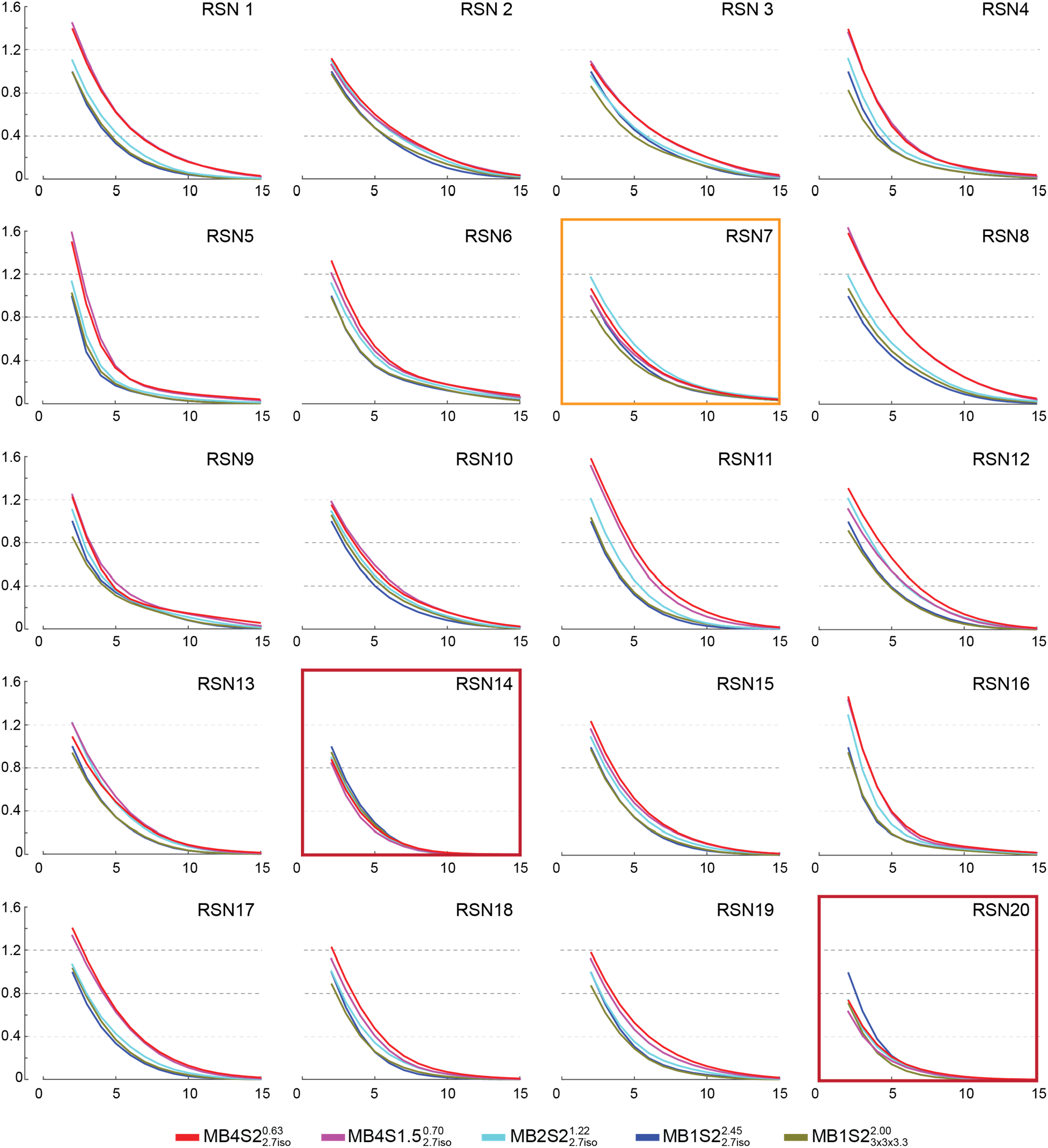
Per component number of voxels surviving any t-threshold relative to the non-accelerated 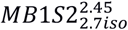 sequence for resting state analysis. The x-axis represents the voxel wise t-values, and the y-axis the number of voxels surviving that threshold relative to the number of voxels surviving *t* ≥ 2 at 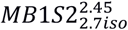. MB4 sequences are most inclusive, except for the plot framed in orange and red, where 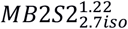 and 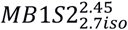 sequences include the largest number of voxels.

## Discussion

We tested if acquisition with a shorter TR and therefore higher sampling rate – as afforded by MB – may improve our power to identify the neural substrates of cognitive functions. Based on our analysis of task-based fMRI data, for group level analyses, the use of MB acceleration is beneficial to improve voxel-wise statistics. In our Philips implementation, sequences with MB4 acceleration (both S1.5 and S2) and 2.7 mm isotropic voxels, showed the highest t-statistics in our task-based fMRI analysis.

Other than the effect of MB acceleration, there were three main factors that influenced the group statistics. First, the impact of voxel size is well known (Robinson, Pripfl, Bauer, & Moser, 2008) and with this data we show that while for tSNR, MB2 is enough to compensate for 50% reduction in voxel volume (see Supplementary analysis, tSNR section, Supplementary Figure 2B), when it comes to the group level t-values, MB4 had t-values that were similar to the t-values seen with MB1 sequence with 3×3×3.3 mm voxel size (Figure 2B,E). Furthermore, higher t-values were observed when the group sizes were bigger (Turner, Paul, Miller, & Barbey, 2018) and the subjects viewed the stimuli only once (Larsson & Smith, 2011). While these are indirect observations from the data presented here, it can be clearly seen from Figure 2 C-E that MB does not compensate for factors such as larger sample sizes and repetition suppression effect.

It should be noted that all the sequences used in the current acquisition were able to identify the core regions that have been shown to be a part of the action observation network (Abdelgabar et al., 2018; Valeria Gazzola & Keysers, 2009). ROC analyses suggest a strong concordance between the outcomes of different sequences (tables 3, 5). So if one selects any of the sequences presented here, they would reach more or less similar conclusion about the brain regions implicated in the cognition of interest. Looking at the areas where we found disagreement between the sequences, we notice that they particularly include subcortical regions including the cerebellum and basal ganglia, which are detected more often in sequences with MB acceleration or with the sequences with bigger voxel sizes (see blue box in table 4). This finding further supports the use of MB acceleration for detecting task correlated BOLD particularly if one is interested in these regions. Moreover, using dual regression analysis we show similar benefits of MB acceleration on RSNs (Figure 5-7) supporting the evidence present in the literature that tests MB acquisition using resting state fMRI (Feinberg et al., 2010; Griffanti et al., 2014; X.-H. Liao et al., 2013).

We looked at inter and intra-subject variability in an attempt to shed light on how MB benefits group analyses. In our previous study where we tested standard single-shot 2D echo planar imaging (EPI) to three advanced EPI sequences, i.e., 2D multi-echo EPI, 3D high resolution EPI and 3D dual-echo fast EPI, inter-subject variability had a major impact in determining the sensitivity of the group-level analyses (Kirilina et al., 2016) in almost all brain regions except of prefrontal cortex, thereby limiting the potential for improving results by improving the measurement of each subject. However, in the current study, we found that the contribution of intra- and inter-subject variance to the random effect group level analysis was comparable. Therefore the dominant form of variance varies across regions, making the distinction of between- and within-subject variance less fruitful in understanding how MB benefits the group analysis (Figure 3).

Looking at other image statistics such as G-factor and tSNR, we see that g-factor values increase with higher MB suggesting increase in noise due to poorer separation of the aliased voxels (Supplementary Figure 1). However, correcting for the number of acquired volumes, the predicted tSNR and the measured effective tSNR (Supplementary Table 2, Figure 2) both improve with higher MB factor. Since the variance shows no systematic differences and the noise measures increase with higher MB, higher sampling rate, increasing the statistical power is the most likely explanation for the better t-statistics with higher MB.

To explore whether the additional volumes acquired with higher MB can compensate for performing fewer repetitions of the task, we truncated the data and examined the first third of each experiment. With one third of the MB4 acquisition containing as many samples as the full MB1 acquisition, one might have expected the former to generate maps quite similar to the latter. However, we see that higher multiband did not show benefits in terms of localizing activations more reliably when fewer repetitions of a condition are available. This is contrary to what has been claimed for rs-fMRI (X.-H. Liao et al., 2013). This inability of the additional data points obtained using multiband to replace those obtained by a longer task is similar to the recent findings showing that temporal down sampling by randomly removing up to 50% of time points has little effects on BOLD reliability, while truncating the datasets is associated with decreased reliability (Shah et al., 2016), and may reflect the fact that temporal autocorrelation in fMRI data may make adjacent data-points acquired with short TR comparatively redundant.

Taking these factors together, our data suggests that on our Philips 3T scanner, and implementation of multiband (based on scanner software release version R5.4), faster scanning offers modest but significant benefits for group-level voxel-wise task-correlated statistics and can be used as a better alternative to the single-band EPI, if other variables such as voxel size, scan duration and sample size are identical. Using MB factor of 4 with in plane SENSE factor of 1.5 or 2 and spatial resolution of 2.7 mm isotropic seems superior to the other sequences used here. However, when deciding whether to invest into multiband technology for cognitive neuroscience paradigms, our analyses suggests that for task-based studies, results similar to MB acceleration of factor 4 can be achieved by increasing the voxel size. This however might not be true for resting state studies. In our resting state analysis, sequence with MB1 and 3×3×3.3mm voxel size showed the lowest t-values. One potential explanation is that while looking at RSN connectivity, smaller voxel size may result in decreased partial volume averaging therefore might improve connectivity estimate compared to the use of larger voxel sizes (Newton, Rogers, Gore, & Morgan, 2012).

One important thing to note here is that our study focuses on comparing group-level statistics. This was chosen, because most cognitive neuroscience studies draw their conclusions from such group studies with ~20 participants. An interesting additional question is whether MB would alter the t-values obtained at the single subject level. This might be particularly interesting when studying rare patients, for instance, or performing pre-surgical scanning. Even though each of our subjects was scanned with all sequences, we found it difficult to answer this seemingly simple question. This is due to two reasons. First, the difference in TR across sequences substantially changes the white-matter/gray-matter contrast (Supplementary Figure 5). As a result, the global mean normalization that is part of the traditional SPM pipeline applies different normalizing factors to images from different TR, and hence makes the beta maps non-comparable. Second, sequences with shorter TR acquire more samples and their single subject t-values thus need to be compared against different critical t-values due to the difference in degrees of freedom. This makes comparing the single subject t-values across sequences meaningless. We tried to compensate for this effect by z-transforming the p values (invnorm(p)) or comparing effect sizes (t/sqrt(df)), but found that doing so leads to counterintuitive results that suggest potential issues with the calculation of degrees of freedom. For a detailed analysis on this topic using the same data presented here, see Bhandari et al., in prep.

Our study has a number of strengths and limitations that should be considered.

### Limitations

Because we only used a block design task, our study cannot address whether event-related designs may benefit more from increased acquisition speed, and this should be investigated in future studies. Also, whether cognitive neuroscience questions which can use MB acquisition without decreasing the TR and thus using the silent periods for presenting auditory stimuli (De Martino et al., 2015) or performing simultaneous EEG recordings (Uji, Wilson, Francis, Mullinger, & Mayhew, 2018) or other electrical recordings that may benefit from reduced interference from the radio-frequency pulses from fMRI, have not been tested here. Next, higher sampling with MB acceleration allows acquisition of high frequency components. Here we do not explore whether these higher frequencies, which are absent in the HRF convolved predictor, contain task-induced neural information. Moreover, shorter TRs offer advantages for spectral de-aliasing (Tong & Frederick, 2014a; Tong et al., 2014). Here we decided not to perform low-pass filtering because studies have shown the presence of BOLD like components in high frequency bands (Boubela et al., 2013). Here we employ 16 regressors including the six motion and ten principal components from CSF and WM (CompCorr, Behzadi et al., 2007) to de-noise the data. However, the effect of other noise cleaning procedures that have been proposed for cleaning MB data such as FIX (Boyacioğlu et al., 2015), have not been tested on this data and may be considered in the future exploration using this data. An additional limitation of our study is that for the MB2 condition, we had 22 instead of 23 participants. Because t-values scale with the square route of the degrees of freedom, this means that we have underestimated the t-values for the MB2 by about 2%. The differences between MB2 and MB4 we report here are however larger than 2%, and we feel that our data therefore nevertheless supports benefits of MB4 over MB2, and performing all analysis on N=22 participants lead to similar results (See supplementary figure 7). Finally, we report that the image grey-white CNR decreases as the TR becomes shorter (Supplementary Figure 3). In this data where the volunteers were healthy adults and there was minimal subject movement, we did not encounter any issues with the realignment steps of pre-processing pipeline. However, in patient population where motion might be an issue, an additional loss of CNR may result in sub-optimal realignment, thereby affecting the final statistics. This concept can be explored using controlled head movements in the scanner and may be addressed in future studies.

### Strengths

While most of the previous studies looked the effect of MB on the summary statistics from the subject level analyses (Demetriou et al., 2018; Kiss et al., 2018; Sahib et al., 2016; Todd et al., 2016), the current study is one of the only 2 studies that look at the effect of MB acceleration on voxel-wise group-level task-related statistics (Boyacioğlu et al., 2015). While the subject-level statics are good indicators of the performance of MB, one should be careful while interpreting the findings coming from sequences that result in variable image CNR as well as have different degrees of freedom per sequence (Bhandari et al., in prep). While most previous studies looked at the effect of multiband on very simple sensory tasks and lenient contrasts, using very small sample sizes (median N = 10), our study used a task and sample size more representative of contemporary cognitive neuroscience studies to have the power to detect even moderate benefits of multiband and ensure that our results are reproducible and representative. That our group results are so similar at different MB and so similar to an independent study (Figure 2) provides evidence for the robustness of our cognitive neuroscience task and results. Using the same participants on the same scanner at different multiband levels ensures that differences in multiband performance are not the result of differences across subject pools or scanners.

In summary, we can thus recommend the use of MB4 on a Philips scanner when aiming to perform group-level analyses using cognitive block design fMRI tasks using voxel sizes in the range of cortical thickness (e.g. 2.7mm isotropic). While results will not be dramatically changed by the use of multiband, our results suggest that MB will bring a moderate but significant benefit.

## Acknowledgements

This work was supported by the Netherlands Organization for Scientific Research (VIDI: 452-14-015 to V.G.), the Brain and Behavior Research Foundation (NARSAD young investigator 22453 to V.G.), the European Research Council of the European Commission (ERC-StG-312511 to C.K.) and the BIAL foundation grant (503 323 055 to V.G., C.K. & R.B.). We thank the Spinoza centre for neuroimaging where the scanning was performed and the staff members of Spinoza centre specially Dr. Pieter Buur and Dr. Wietske van der Zwaag for their input on the experimental protocol and comments on the manuscript.

## Supplementary Materials

**Supplementary table 1.** List of the hand actions used in action observation. The actor’s right hand always entered the screen and initiated the goal-directed action from the left side of the screen. Note that control movies showed the same objects used for action observation but the actor’s hand moved close to but without interacting with the object.

| N° | Recorded actions description | Movie Duration |
| --- | --- | --- |
| 1 | Stirring coffee with a spoon. | 2s |
| 2 | Putting a cube of sugar in a cup of coffee. | 2s |
| 3 | Closing a box with a key. | 2s |
| 4 | Lighting a candle with a lighter. | 2s |
| 5 | Putting a flower in a vase | 2s |
| 6 | Putting a battery in a remote control. | 2s |
| 7 | Putting a CD in a CDs stack. | 3s |
| 8 | Hammering a nail. | 3s |
| 9 | Putting whipped cream on strawberries. | 3s |
| 10 | Cutting a deck of cards. | 2s |
| 11 | Placing jewelry in a box. | 2s |
| 12 | Crumpling a paper sheet. | 2s |
| 13 | Closing a box of chewing gums. | 2s |
| 14 | Putting a pin on a foam base. | 2s |
| 15 | Taking hand cream from a tin. | 2s |
| 16 | Taking some tape and placing it on a box. | 3s |
| 17 | Pouring wine in a glass | 3s |
| 18 | Watering a plant. | 3s |
| 19 | Stirring eggs. | 2s |
| 20 | Closing a water bottle. | 2s |
| 21 | Flipping through a block note. | 2s |
| 22 | Taking an olive from a jar. | 2s |
| 23 | Putting a candle in a candleholder. | 2s |
| 24 | Closing a folder. | 2s |
| 25 | Cracking walnuts. | 3s |
| 26 | Placing a wine bottle in a box. | 3s |
| 27 | Opening a suitcase. | 3s |
| 28 | Spreading jam on a piece of bread. | 2s |
| 29 | Cutting a ribbon on a package. | 2s |
| 30 | Stirring soup with a spoon. | 2s |
| 31 | Putting business cards in a box. | 2s |
| 32 | Putting a hair clip in a purse. | 2s |
| 33 | Disconnecting headphones from an MP3 player. | 2s |
| 34 | Breaking an egg on the edge of a bowl. | 3s |
| 35 | Stirring a painting brush in a cup of water. | 3s |
| 36 | Taking a walnut with chopsticks and placing it in a box. | 3s |

### Supplementary Analysis 1: G-factor

To get an estimate of the aliasing noise that results from sub-optimal separation of the simultaneously acquired voxels, g-factor maps were inspected. G-factor values closer to 1 are considered “clean” and higher values in the voxels represent more noise. Supplementary figure 1 shows g-factor maps from a representative subject for 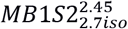, 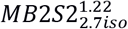, 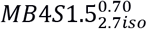. The histogram of these maps is shown in supplementary figure 2. As expected, the aliasing noise increases with higher MB factors. While the noise amplification in 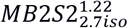 is restricted to a small region (hot spots in the posterior part of the images, row 2), the sequences with 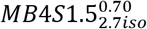 show more widespread g-factors above 1.2 (yellow in top row). Row 3 in supplementary figure 1 shows the g-factor values for 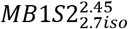 and the values fall much closer to 1 - as expected, since there are no simultaneously acquired voxels. Table 3 presents the average g-factor mode and the 99 percentile values over all voxels and subjects. As in the single subject maps, at the group level we can see an increase in g-factor mode from MB1 to MB4. To understand the effect of g-factor penalty on tSNR, we calculated the relative expected tSNR (tSNR_rel_). Practically, tSNR_rel_ was computed per voxel relative to our reference sequence with given TR_ref_=2.45 and SENSE factor S_ref_=2 without MB implementation using equation s1.

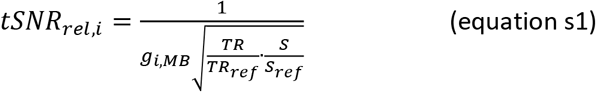

Here, g_i,MB_ is the g-factor value in voxel i, for a certain sequence. Supplementary table 2 presents the tSNR_rel_ mode and the 99 percentile values over all subjects. We find that despite higher g values, the predicted tSNR_rel_ increases in sequences with higher MB factor. While the higher g-factor values speak against the use of higher MB factors, in theory, reduction in TR leading to more data points per unit time, would ultimately result in an 87% (√(2.45/0.70)-1) increase in effective SNR going from MB1 to MB4. This can be seen to be true in tSNR_rel_ values. Thus, if we base our conclusion on these matrices, scanning at higher MB-factors is preferable.

**Supplementary Figure 1.**
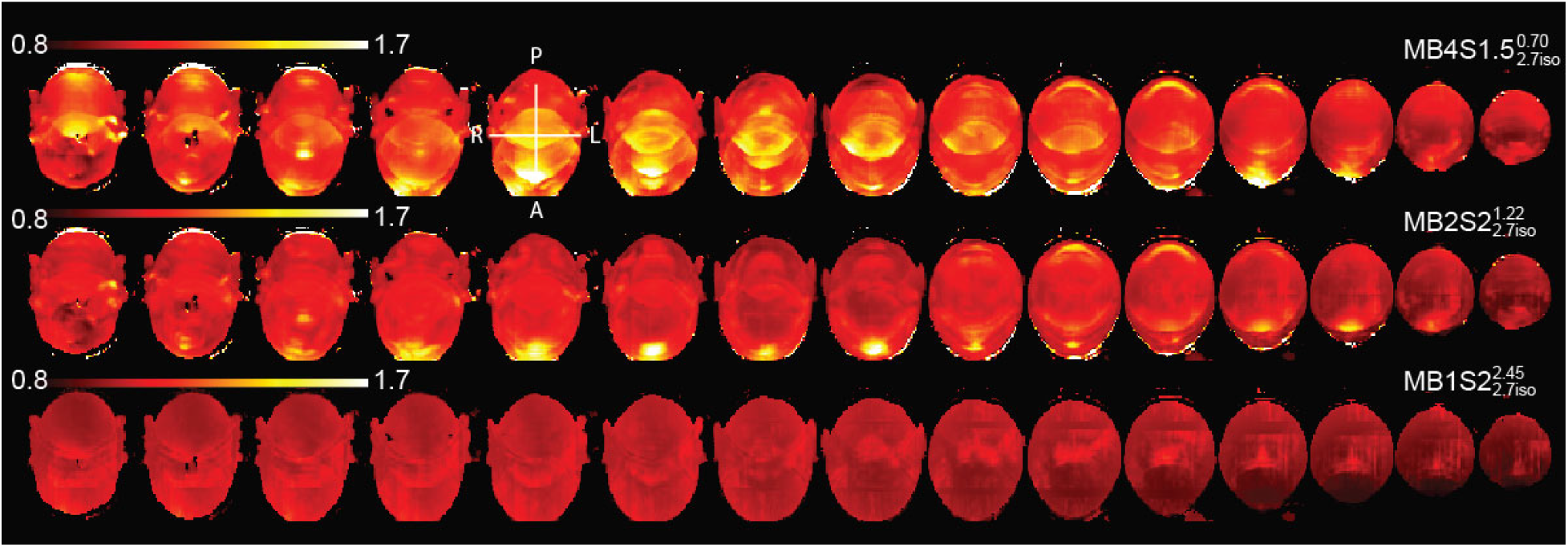
Raw G-factor images from a representative subject for sequences 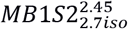, 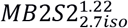 and 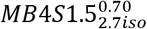. G-factor values close to 1 (red) represent minimal aliasing noise while increase in this value (yellow) indicate suboptimal separation of simultaneously acquired voxels. High values are rare at TR 2.45s, restricted in TR 1.22s but widespread in TR 0.70s.

**Supplementary Figure 2.**
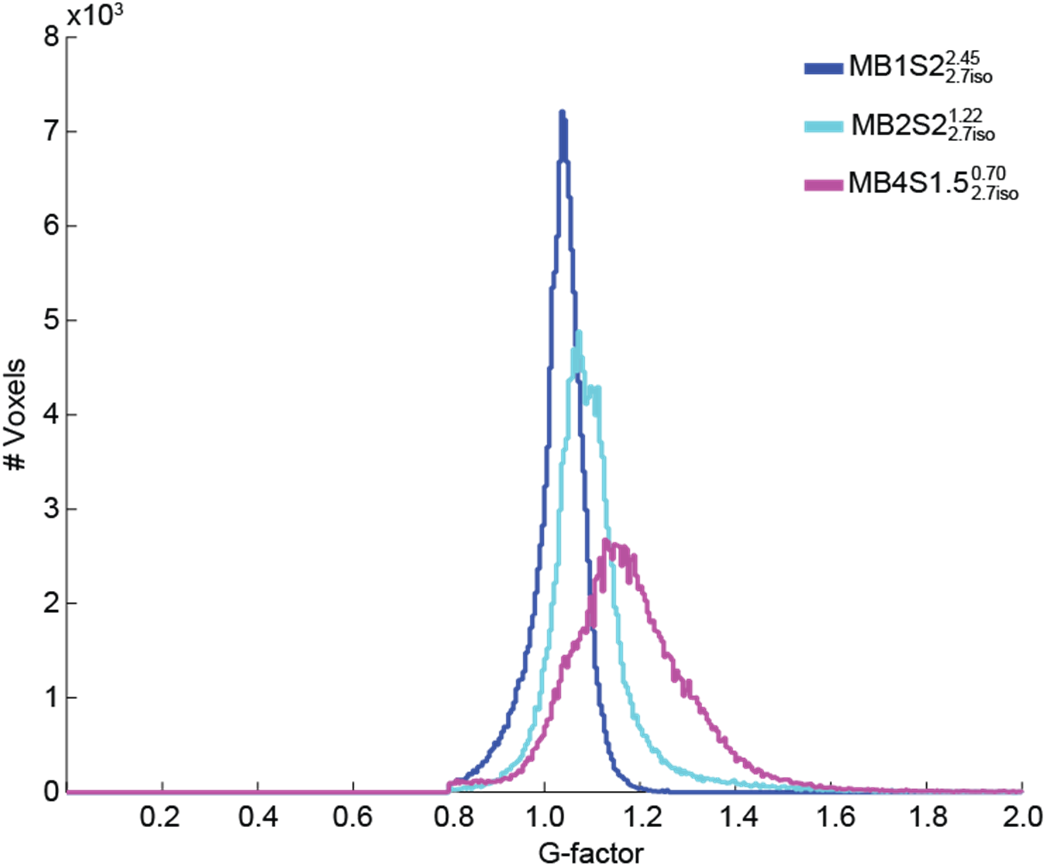
Histogram of images in Supplementary figure 1 from a representative subject confirm that the g-factor values increase when higher MB factor is implemented.

**Supplementary table 2.** G-factor and relative tSNR (predicted)

| TR | | $MB1S2^{2.45}_{2.7iso}$ | $MB2S2^{1.22}_{2.7iso}$ | $MB4S1.5^{0.70}_{2.7iso}$ |
| --- | --- | --- | --- | --- |
| metric | statistic | Mean ± SD over subjects | | |
| G-factor | mode | $1.04 \pm 0.01$ | $1.09 \pm 0.02$ | $1.17 \pm 0.03$ |
| | 99 percentile | $1.14 \pm 0.02$ | $1.52 \pm 0.07$ | $1.65 \pm 0.05$ |
| tSNR<sub>rel</sub> | mode | $0.96 \pm 0.01$ | $1.30 \pm 0.02$ | $1.85 \pm 0.04$ |
| | 99 percentile | $0.88 \pm 0.02$ | $0.93 \pm 0.04$ | $1.31 \pm 0.04$ |

### Supplementary Analysis 2: Effective Temporal Signal to Noise Ratio (effective tSNR)

While the tSNR_rel_ gives a theoretical prediction of the tSNR that can be expected based on data from the initial calibration, we next used the fMRI data to get the actual effective voxel-wise tSNR as described in Todd et al., 2016. We divided the mean signal over time after removal of the task based activity (i.e. m; the constant term from the first level GLM fit) by the standard deviation (σ) over time of the residual signal after the GLM fit. We then scaled this by a factor that corrects for the different number of volumes acquired with different sequences and accounts for the autocorrelations in the data using equation s2.

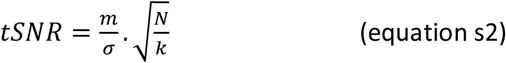

Here, N = number of volumes, k = c^T’^ (X^T’^X) c, where, X = whitened and high pass filtered design matrix and contrast (c) = [0 0 0 0 0 0 0 0 0 0 0 0 0 0 0 0 0 0 1] (i.e. only considering the global factor that captures the average activity after task removal).

The tSNR maps per subject were then masked with a grey matter mask and then averaged across subjects, separately for each acquisition sequence. Supplementary figure 3 presents the average tSNR maps. Visual comparison of the maps shows that the tSNR values are higher with higher acceleration. A within-subject ANOVA with the mean tSNR value in the gray-matter per subject showed a main effect of TR (*F* (4, 92) = 8.61, *p*<0.001) (Supplementary figure 4). Keeping voxel size constant, tSNR values increase with higher MB 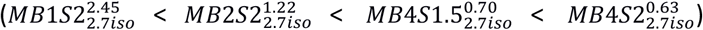. Comparing these values against 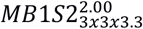 sequence shows that MB2 can compensate for a 50% reduction in voxel volume, and that MB4 provides highest tSNR.

**Supplementary Figure 3.**
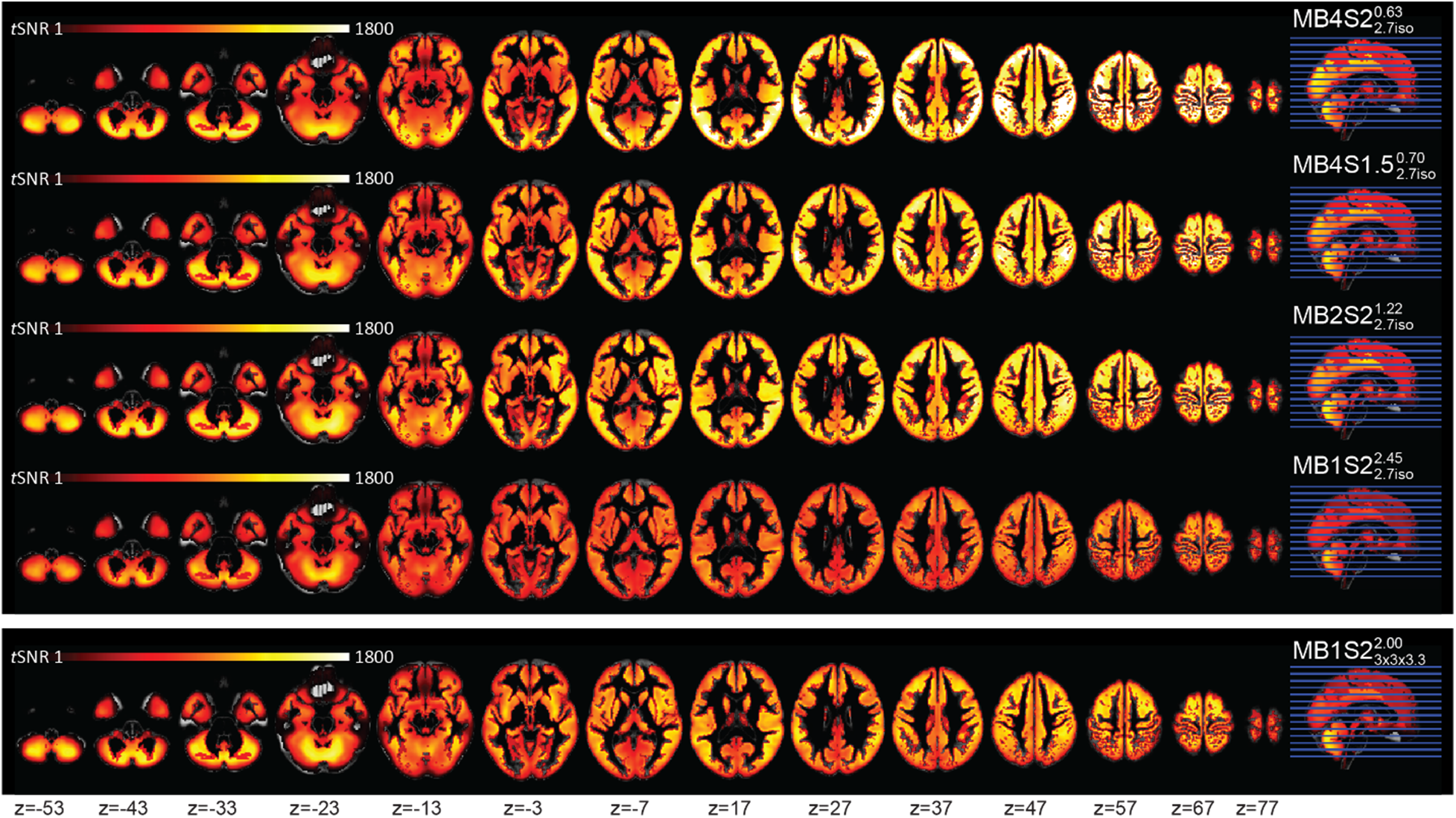
Average TSNR maps (within grey matter mask) as a function of TR. In general, tSNR values appear to be higher for sequences with MB implementation.

**Supplementary Figure 4.**
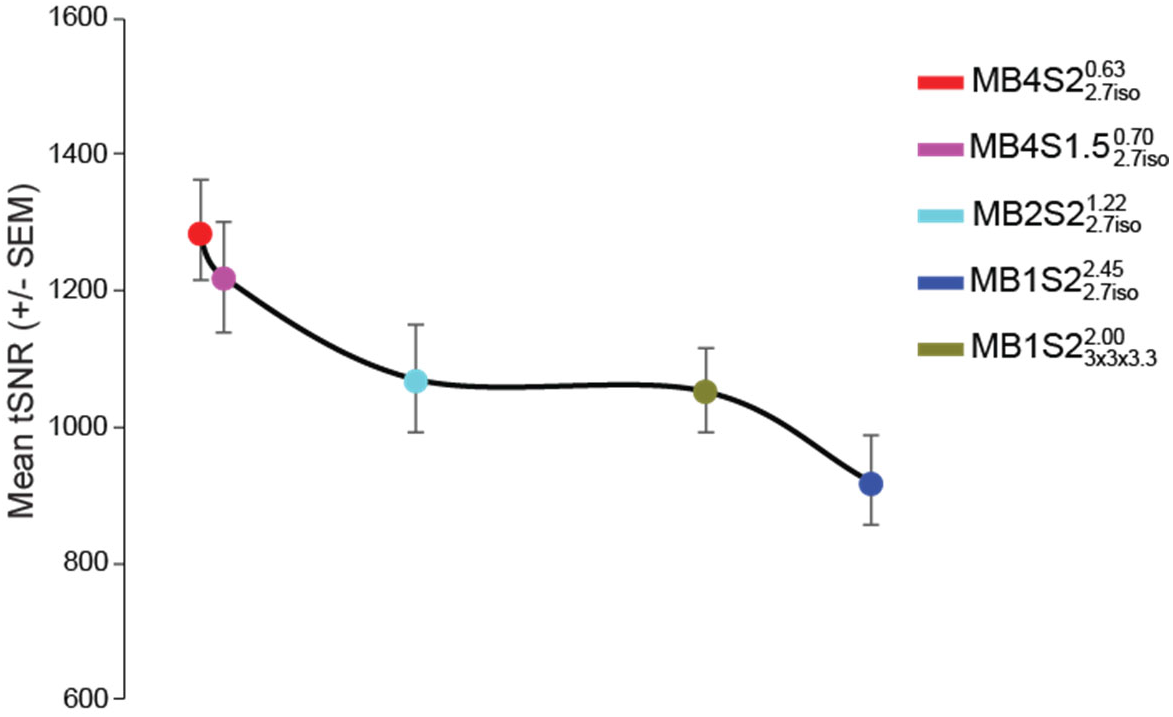
Plot of the mean tSNR values (+/- sem over the subjects) within the grey matter mask confirms that values are higher with higher MB. 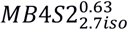 shows highest tSNR values.

### Supplementary Analysis 3: Contrast to Noise Ratio (CNR)

Supplementary figure 5 shows the mean functional EPI as a function of sequence from one representative participant. As can be seen, the white/gray matter contrast is lower for sequences with smaller TRs. For each subject, to quantify this white/gray matter contrast loss, CNR was calculated on the mean functional image (created during the realigning procedure). The mean functional image was separated in white and gray compartments using individual gray and white matter masks obtained from the segmentation of the co-registered T1 image (thresholded to include values > 0.8 of estimated probability). We then calculated the average and standard deviation separately for the gray and white matter compartments. The contrast to noise ratio was then calculated by dividing the signal (i.e. the difference in mean between the white and gray matter compartments) by the noise (i.e. the standard deviation of the union of the demeaned gray and white matter voxels) as in equation s3.

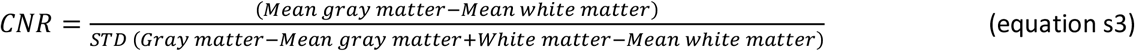

A within-subject ANOVA to test the difference between the CNR values of the five acquisition schemes revealed a highly significant main effect of MB (*F* (4, 84) =512.53, *p* < 0.001) on CNR due to a decrease in CNR with higher TR (Supplementary figure 6). While the decrease was mostly linear with decreasing TR, the CNR values for 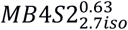 were slightly better than 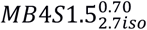, suggesting that in plane acceleration of SENSE 2 may afford slightly better grey-white contrast than SENSE 1.5.

**Supplementary Figure 5.**
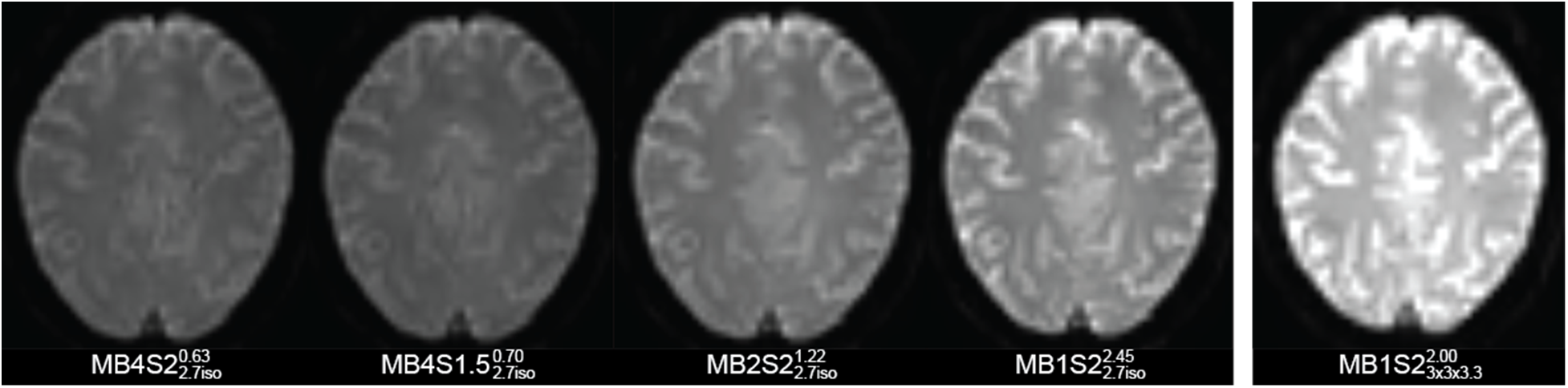
Mean EPIs from a representative subject estimated during the realignment procedure. The images are not normalized and they correspond to z=31. The grey white contrast seems to decrease for smaller TRs and artifacts become evident for MB>2.

**Supplementary figure 6.**
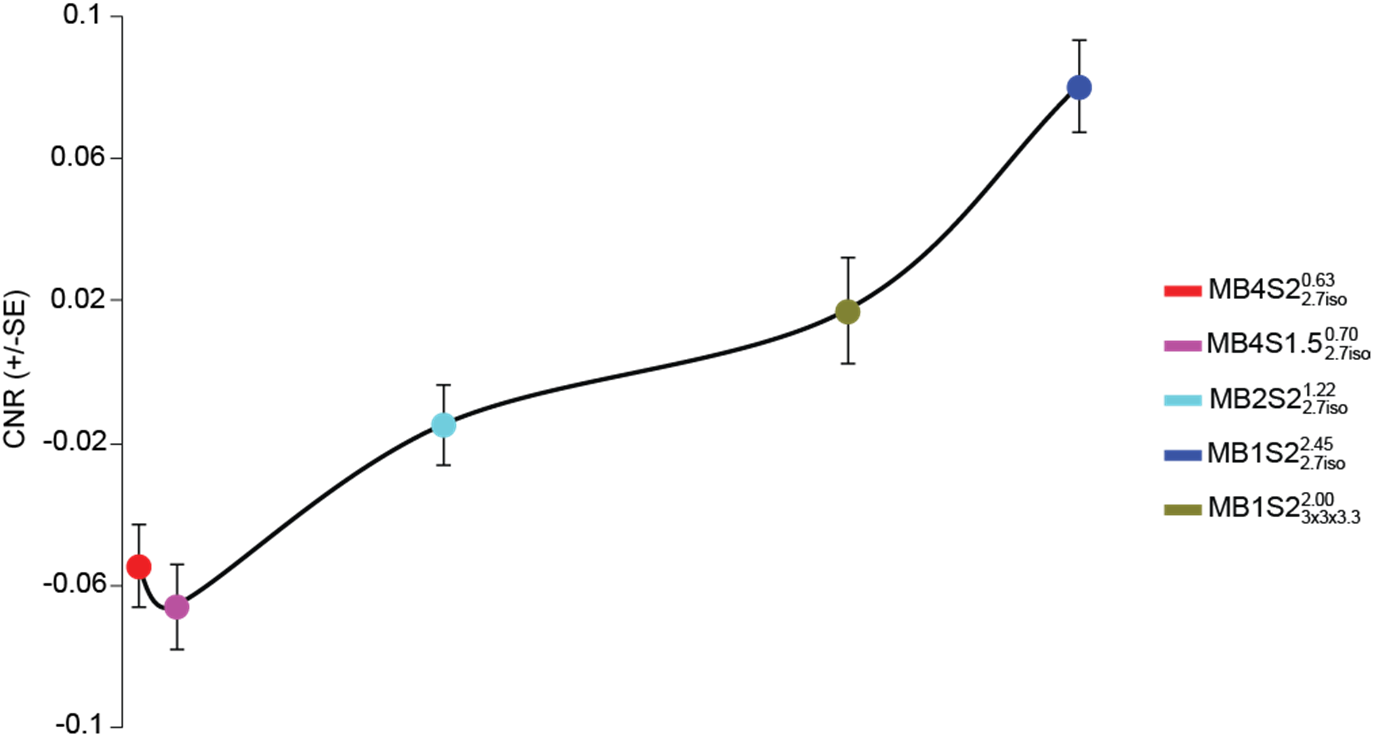
CNR of the white/gray matter shows a significant main effect of MB (*F*(4, 84) = 512.5, *p*< 0.001) confirming a decrease in contrast values as the TR becomes smaller.

### Second-level t-test with 22 subjects

Second level t-tests were performed with 22 subjects for each sequence, making the sample size equal to that with MB2 acceleration. Histograms of the t-values from these t-tests are presented in supplementary figure 7. It is apparent that that the overall results and conclusions do not chance when a sample of 22 vs 23 is used. See figure 2 D for comparison.

**Supplementary figure 7.**
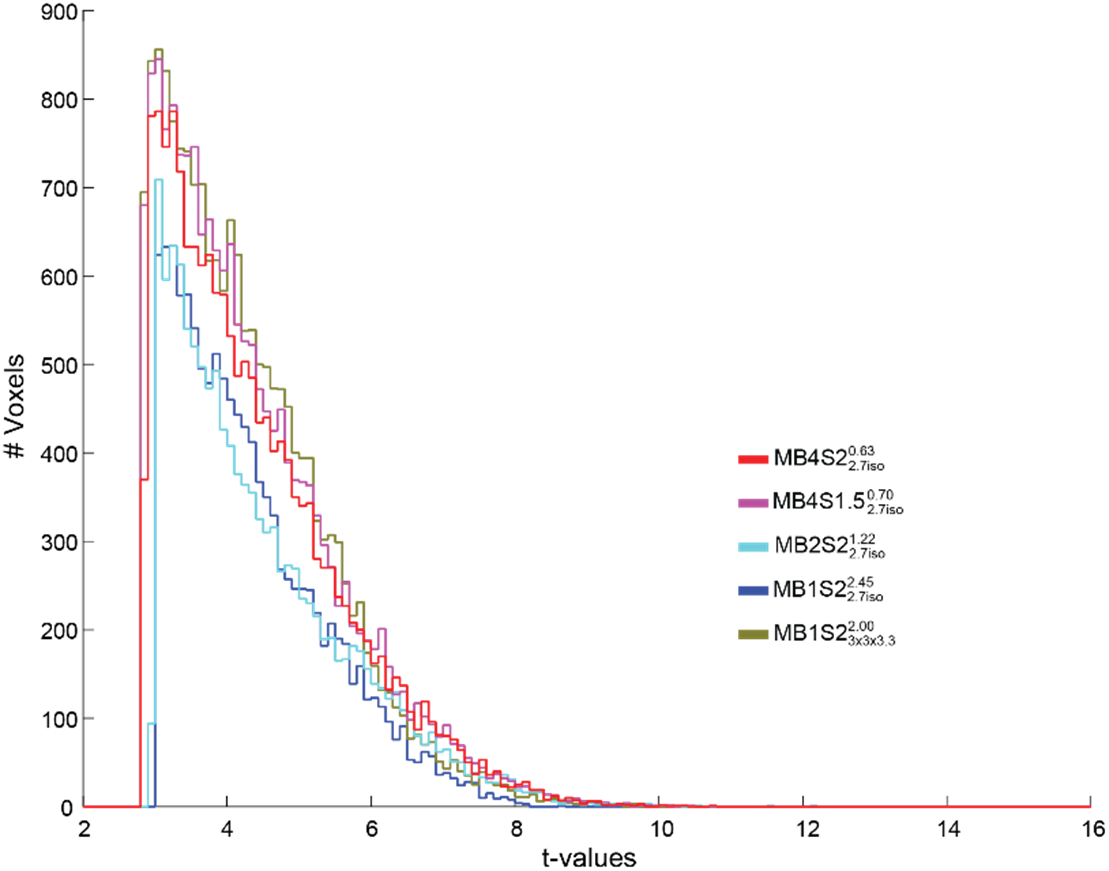
If we use 22 subjects for each TR, making the sample size same in all sequences, the overall t-values and their distribution between different sequences does not change.

